# Single-cell analysis identifies distinct macrophage phenotypes associated with pro-disease and pro-resolving functions in the endometriotic niche

**DOI:** 10.1101/2024.03.07.583861

**Authors:** Yasmin Henlon, Kavita Panir, Iona McIntyre, Chloe Hogg, Priya Dhami, Antonia O. Cuff, Anna Senior, Niky Moolchandani-Adwani, Elise T. Courtois, Andrew W Horne, Matthew Rosser, Sascha Ott, Erin Greaves

**Affiliations:** Division of Biomedical Sciences, Warwick Medical School, University of Warwick, Coventry, United Kingdom (UK); Centre for Early Life, University of Warwick, Coventry, UK; Centre for Reproductive Health, Institute of Regeneration and Repair, The University of Edinburgh, Edinburgh, UK; Single Cell Biology Lab, The Jackson Laboratory for Genomic Medicine, Farmington (CT), USA

**Keywords:** Lesion, phenotype, heterogeneity, endometriosis, macrophage

## Abstract

Endometriosis negatively impacts the health-related quality of life of 190 million women worldwide. Novel advances in non-hormonal treatments for this debilitating condition are desperately needed. Macrophages play a vital role in the pathophysiology of endometriosis and represent a promising therapeutic target. In the current study, we revealed the full transcriptomic complexity of endometriosis-associated macrophage subpopulations using single-cell analyses in a preclinical mouse model of experimental endometriosis. We have identified two key lesion-resident populations that resemble i) tumour-associated macrophages (characterized by expression of *Folr2*, *Mrc1*, *Gas6* and *Ccl8+*) that promoted expression of *Col1a1* and *Tgfb1* in human endometrial stromal cells and increased angiogenic meshes in human umbilical vein endothelial cells, and ii) scar-associated macrophages (*Mmp12, Cd9, Spp1, Trem2*+) that exhibited a phenotype associated with fibrosis and matrix remodelling. We also described a population of pro-resolving large peritoneal macrophages (LpM) that align with a lipid-associated macrophage phenotype (*Apoe, Saa3, Pid1*) concomitant with altered lipid metabolism and cholesterol efflux. Gain of function experiments using an Apoe mimetic resulted in decreased lesion size and fibrosis, and modification of peritoneal macrophage populations in the preclinical model. Using cross-species analysis of mouse and human single-cell datasets, we determined the concordance of peritoneal and lesion-resident macrophage subpopulations, identifying key similarities and differences in transcriptomic phenotypes. Ultimately, we envisage that these findings will inform the design and use of specific macrophage-targeted therapies and open new avenues for the treatment of endometriosis.

## Introduction

Resident tissue macrophages are integral to the maintenance of healthy tissue function. They exhibit heterogeneity in phenotype and function, and their roles within tissues are dictated by ontogeny, local environment, inflammation status, and time in residence within the microenvironment. These modifying factors mean that each adult tissue contains a unique balance of ontogenetically distinct macrophage populations, and the complement is dynamically modulated throughout life^1^. Tissue-resident macrophages in different tissues and at different time-points arise from three different origins: early yolk-sac macrophages, foetal liver monocytes, or bone-marrow-derived monocytes. In some tissues, macrophages of embryonic origin can exist independently of monocyte input and are maintained locally. In other tissues, monocytes continually enter the tissue and replenish macrophage populations^2, 3, 4^. Inflammatory challenge / injury results in rapid recruitment of monocytes to damaged tissues and a disruption of tissue macrophage homeostasis^1^. Peritoneal cavity macrophages play a vital role in immune surveillance of the cavity and visceral organs, and are an exemplar of the dynamic mosaic of macrophages in tissues. Two main populations of macrophages exist in the cavity: large (LpM) and small (SpM) peritoneal macrophages. LpM are tissue-resident, abundant and predominantly embryonically-derived in early life. In adulthood, LpM are gradually replenished by monocytes in a sexually dimorphic pattern^5^. Although monocyte-derived LpM acquire characteristics of embryo-derived LpM, they exhibit some transcriptional and functional differences^6, 7^. Conversely, SpM are less abundant, consist of monocyte-derived macrophages, and dendritic cells (DC) and are constantly replenished from infiltrating monocytes^5^. These steady-state dynamics are rapidly perturbed by inflammation, that can lead to recruitment of large numbers of inflammatory macrophages and a loss of LpM^8, 9, 10^, the extent of which varies depending on the inflammatory stimuli^7^. In parallel to their role in cavity homeostasis, peritoneal macrophages are also central to peritoneal pathologies including endometriosis^11, 12^.

Endometriosis is an incurable inflammatory condition characterized by the growth of endometrial-like tissue as ‘lesions’ outside the uterus, usually on the lining of the peritoneal cavity or ovaries^13^. The condition is associated with chronic pelvic pain and infertility and affects approximately 190 million people worldwide^14^. Removal of lesions during laparoscopic surgery can relieve symptoms temporarily but recurrence rates are high. Current medical management is via ovarian suppression, which is contraceptive, and symptoms return following cessation of treatment^14^. There is an urgent unmet need for new therapeutic targets that may be developed into novel non-invasive, non-hormonal treatments. Disease-modified macrophages represent a promising focus for the development of immune therapies for endometriosis as they become adapted such that they support lesion survival by promoting cell proliferation and vascularization^15^. Using our unique syngeneic mouse model of experimental endometriosis^16^, we have also demonstrated that macrophages encourage innervation of lesions, sensitization of nerves and generation of pain^17, 18^.

Recently, we identified that macrophages in lesions have different origins and associated functions; they are derived from the eutopic (donor) endometrium, from (host-derived) LpM that traffic into lesions, and monocytes that infiltrate lesions and differentiate into macrophages^19^. Endometriosis also triggers continuous recruitment of monocytes to the peritoneal cavity and leads to heightened monocyte input into the LpM pool. Using genetic and pharmacological depletion strategies we demonstrated a ‘pro-endometriosis’ role for endometrial macrophages and an ‘anti-endometriosis’ role for monocyte-derived LpM^19^. Although the complexity of macrophage origin and function in endometriosis is now partially revealed, the true complexity of macrophage phenotype, and how disease modified macrophages differ from macrophages in steady-state tissues is currently unknown. The identification of unique markers that discriminate ‘pro-endometriosis’ macrophages, will facilitate a targeted therapeutic approach to eliminate or alter disease-promoting macrophages, whilst leaving those critical for normal physiological tissue function intact. In the current study, we have used single cell transcriptomics (scRNA-Seq) to investigate the transcriptional heterogeneity of both lesion-resident and associated peritoneal macrophages, and to identify population specific markers. We have further explored the role of specific phenotypes using *in vitro* and *in vivo* functional studies, shedding new light on the role of macrophages in the pathophysiology of endometriosis. Additionally, we performed cross-species mapping of human and mouse datasets to determine the concordance of populations between the species. Taken together, we describe critical background knowledge required for the development of future targeted immunotherapy for endometriosis.

## Results

### Endometriosis-associated macrophages exhibit phenotypic heterogeneity and unique markers

To further understand the dynamics of endometriosis-associated macrophage subpopulations and to identify phenotypic heterogeneity we performed 10X scRNA-Seq on isolated leukocytes from samples recovered from a mouse model of experimental endometriosis^16^. In brief, endometriosis was induced in wild-type C57BL/6 recipient mice (recipients were ovariectomised and supplemented with oestradiol) by injecting ‘menses’ like donor endometrium into the peritoneal cavity and allowing lesions to establish over two weeks (Endo-Ovx model). We isolated CD45+ leukocytes from lesions and the peritoneal lavage of mice with endometriosis using FACS. For comparison, we also isolated leukocytes from sham mice (mice were ovariectomized, supplemented with oestradiol and subject to an injection of intra-peritoneal saline instead of endometrial tissue) and eutopic ‘menses’-like endometrium (4 hours post progesterone withdrawal) from ‘donor’ mice. An aggregate analysis of CD45+ peritoneal lavage cells, ‘menses’ endometrial tissue and endometriosis lesions was performed (Fig.1A). UMAP projection revealed 18 clusters (Fig.1B). We ascertained cell identity of each cluster using canonical markers (Fig.S1); 13 of these were monocytes / macrophages / DCs based on expression of *Ccr2*, *Adgre1, Cd209a,* or *H2-Aa* expression. Other clusters were B cells, T cells, and NKs. UMAP projection based on sample ID revealed that a proportion of lesion-resident macrophages clustered with LpM (arrow) and endometrial macrophages (rings; Fig.1B inset) highlighting our previous observations that lesion-resident macrophages have different origins^19^.

**Figure 1:**
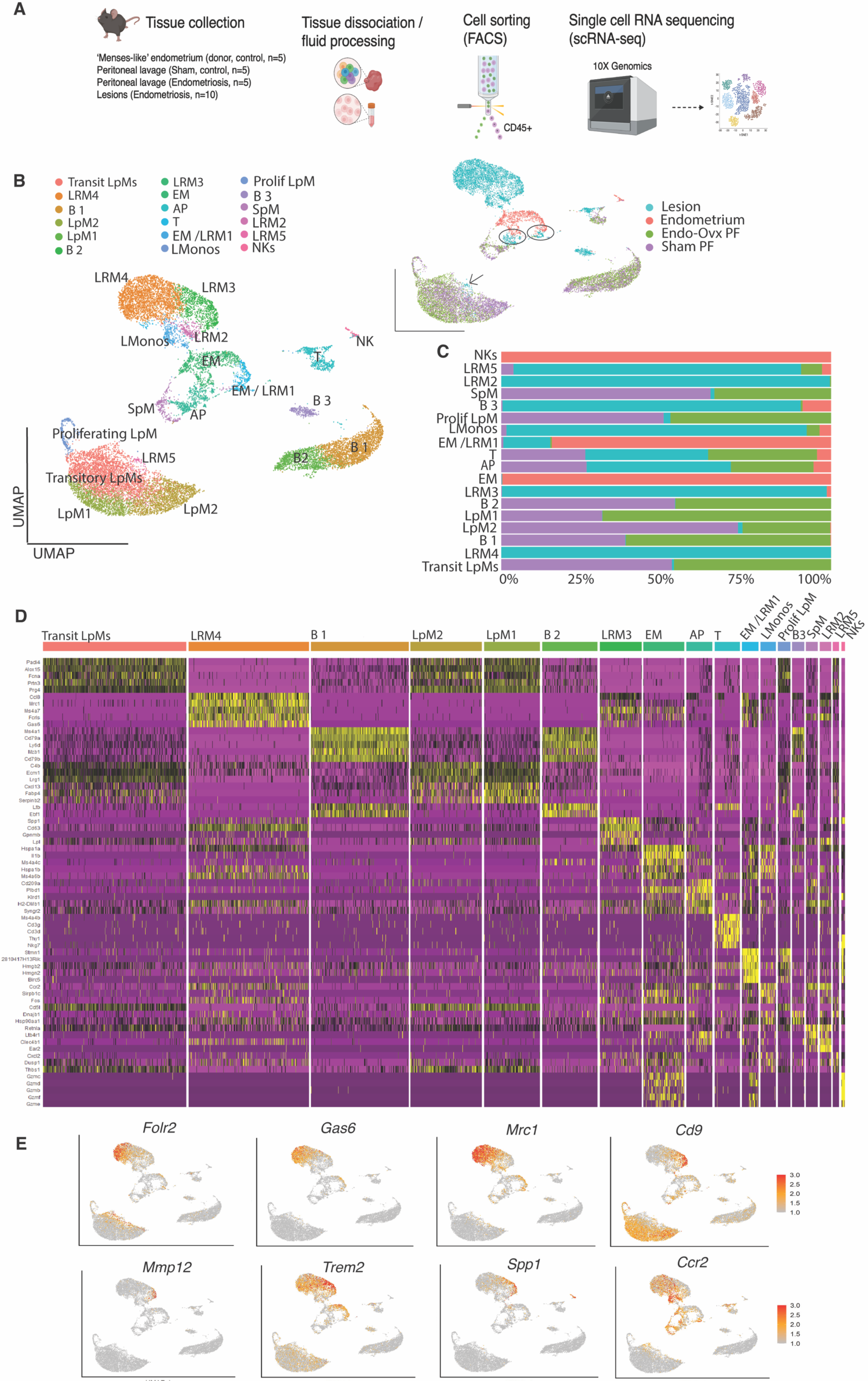
Endometriosis-associated macrophages exhibit significant transcriptomic heterogeneity. A) Schematic of workflow and tissues, fluid evaluated. B) UMAP projection of CD45+ cells isolated from ‘menses’-like endometrium (donor mice; n=5), peritoneal lavage (PF) from sham mice (n=5), PF (n=5) and lesions (n=10 mice) isolated from mice with endometriosis. Inset is UMAP projection based on library ID. AP; antigen presenting cells, EM; endometrial macrophages; LpM; large peritoneal macrophages, LRM; lesion resident macrophages, SpM; small peritoneal macrophages, LMonos; lesion resident monocytes, NK; natural killer cells. C) Bar chart depicting cluster membership of each sample type. D) Heatmap of top 5 differentially regulated genes (DEGs) for each cluster. E) Feature plot of marker genes exhibiting restricted expression in SAM-like and TAM-like lesion-resident macrophages.

Endometrial-derived macrophages separated into 3 clusters, 2 of which were shared by lesion-resident macrophages. The most abundant endometrial cluster (EM) was characterized by the differentially expressed genes (DEGs) *Il1b*, *Ly6c2*, *Ms4a4c*, *S100a9* and *S100a8* (Fig.1D; see SI Table 1 for gene list). Expression of *Ly6c2* and *Ms4a4c* indicate that this population was composed predominantly of monocytes. The second cluster (AP) containing endometrial cells was characterized by the DEGs *Cd209a*, *Slamf7, Cd74* and *Lsp1,* suggesting a mixed population of antigen presenting cells (including DC) and was comprised of endometrial, peritonea, and lesion-resident cells. The final endometrial cluster (EM / LRM1) was characterized by expression of components of the cell cycle pathway highlighting that proliferating macrophages are present in both ‘menses-like’ endometrium and lesions.

Peritoneal macrophages separated into 5 clusters comprising 1 cluster of SpM (*H2-Aa*, *Ccr2*, *Retnla*) and 4 clusters of LpM (LpM1, LpM2, Proliferating LpM and Transitory LpM). Proliferating LpM were identified based on their expression of cell cycle / proliferative markers such as *Top2a, Cdk1, Mcm5, Mki67* and were represented equally in both Endo-Ovx and Sham samples (Fig.1C). Both Endo-Ovx and Sham mice exhibited a predominant population of ‘transitory’ LpM (Fig.1B-C, see also Fig.S2) which were characterized by a down-regulation of *Mrc1*, *Ccr2* and *H2-Aa* and the emergence of LpM markers including *Icam2.* In support of these cells being ‘transitory’, GO and KEGG analysis (SI Table 2) did not return any unique terms for this cluster, with many terms primarily related to translation; in line with macrophages responding to their environment and transitioning through functional states. The presence of a large population of transitory LpM corroborates data in previous studies indicating that surgery to remove the ovaries (with or without oestradiol supplementation) significantly impacts the peritoneal immune environment in female mice resulting in increased macrophage replenishment^6^. To begin to differentiate between embryo-derived / long-lived tissue-resident LpM and monocyte-derived LpM we initially evaluated the presence of *Ccr2* (marker of recently recruited monocytes /monocyte-derived macrophages) and *Timd4* (long-lived tissue-resident macrophages). *Ccr2* was most abundant on SpM, with expression on some transitory LpM. *Timd4* was most prominent on LpM1 and LpM2 with some sporadic *Timd4*+ cells in the transitory population (Fig.S2D). LpM 1 were characterized by DEGs shared by steady-state prototypical LpM: *Cxcl13, Prg4, Alox15*, and *Fabp4* and the emergence of *Tgfb^6^*. Evaluation of cluster occupancy revealed shifts in LpM populations; prototypical LpM (LpM 1) were expanded, whilst LpM 2 (characterised by *Fcna, C4b, Ecm1,* and *Lrg1* mRNAs) were depleted in Endo-Ovx mice (Fig.1B and C).

Lesion-resident macrophages separated into an additional 5 clusters. LMono expressed *Ccr2* and *Ly6c* as well as *Il1b,* indicating the cluster is comprised predominantly of monocytes. LRM2 DEGs were those traditionally associated with monocyte-to-macrophage transition and monocyte-derived inflammatory cells such as *Ccr2*, *Ear2,* and *Clec4b1*^20^. Unique GO terms included those associated with ‘dendrite morphogenesis’ and ‘regulation of synapse activity’, the KEGG pathways returned were ‘prion disease’, ‘Alzheimer’s disease’, and ‘pathways of neurodegeneration’ (SI Table 2) indicating that this subpopulation may be associated with pathways of neurogenesis and neuroinflammation and supports studies (including our own), that endometriosis-associated macrophages are implicated in pathogenic recruitment and activation of nerves^17, 21^. Other GO terms included ‘negative regulation of lymphocyte differentiation’, ‘negative regulation of T cell differentiation’, and ‘negative regulation of leukocyte differentiation’. These results indicate a role for this population in the suppression of specific immune cell differentiation pathways as a mechanism to modulate immune response in the endometriotic environment, perhaps by contributing to immunotolerance of lesions. LRM3 exhibited a transcriptional profile shared with scar-associated macrophages (SAMs) / fibrosis-associated macrophages^22^ characterized by the pro-fibrogenic genes *Spp1* and *Lgals* as well as *Trem2*, *Mmp12*, *Cd9* and *Gpnmb.* Using feature plots, genes exhibiting cluster-restricted expression were visualized (Fig.1D). *Mmp12* and *Spp1* exhibited expression specific to SAM-like cells, whereas *Cd9* and *Trem2* mRNAs were most abundant in SAM-like cells but also exhibited expression, albeit at lower levels, in some other clusters. Unique terms returned for LRM3 included ‘ensheathment of neurons’, ‘ensheathment of axons’, ‘neuron migration’ as well as ‘CD4-positive, alpha-beta T cell activation’ and ‘negative regulation of tumour necrosis factor production’, indicating that this subpopulation is involved with neuron migration, promoting T-cell response, and limiting production of pro-inflammatory cytokines, all supporting these macrophages as exhibiting a pro-repair phenotype. Of the KEGG pathways returned, ‘proteoglycans in cancer’ was of particular interest given the function of proteoglycans as key components of extra-cellular matrix (ECM) that influence cell migration, invasion, and angiogenesis^23^ and indicates that these macrophages may play an important role in ECM formation, remodelling and fibrosis. The most abundant lesion-resident population was LRM4 which was characterized by expression of *Ccl8*, *Mrc1*, *Gas6*, *Marks, Cbr,* and *Folr2,* a signature shared with many tumour-associated macrophages (TAM)^24, 25, 26^. Other interesting genes that were differentially expressed in this population included *Sepp1*, *Nrp1,* and *Igf1*. Visualisation using feature plots demonstrated that *Gas6* and *Folr2* exhibited predominantly restricted expression to TAM-like cells, whereas *Mrc1* exhibited lower expression in some other clusters (Fig.1E). GO terms returned for TAM-like macrophages included ‘central nervous system development’, ‘positive regulation of nervous system development’, ‘mesenchymal cell differentiation’, ‘fibroblast migration’, ‘vasculogenesis’, and ‘response to ischemia’ consistent with a role for these macrophages in neuroangiogenesis, tissue remodelling and repair and response to hypoxic conditions. KEGG pathways also included ‘neuroactive ligand-receptor interaction’. Finally, LRM5 clustered with peritoneal macrophage populations and likely represents peritoneal macrophages that have recently infiltrated into lesions. LRM5 shared several genes with LpM such as *Prg4* and *Saa3* (Fig.1D). Unique GO terms included ‘response to pain’, ‘response to fibroblast growth factor’, ‘keratinocyte proliferation’ and ‘smooth muscle cell differentiation’ highlighting potential roles in tissue repair and remodelling, cellular crosstalk, and pain modulation. *Ccr2* was also visualised using a feature plot (Fig.1D) and exhibited highest expression in LMonos and LRM2 with lower expression extending out into LRM4, EM and AP.

### SAM-like and TAM-like cells arise from monocyte precursors and not infiltrating peritoneal macrophages

We previously determined that macrophages in endometriosis lesions have different origins^19^. In this study, we sought to align lesion-resident macrophage origin to the different phenotypes uncovered using single-cell discovery. To achieve this, we performed reciprocal transfer and adoptive transfer experiments using cells derived from MacGreen (Csf1r-EGFP) mice (Fig.2A) to facilitated isolation of ontogenetically distinct populations from lesions followed by qPCR for TAM-like and SAM-like markers. To isolate endometrial-derived monocytes / macrophages from endometriosis lesions we injected ‘menses’-like endometrium from MacGreen donors into the peritoneal cavity of wild-type recipients. At 2-weeks post tissue-injection lesions were collected, digested and FACS sorted into GFP+ (endometrial-derived) and GFP-(recipient-derived) monocytes / macrophages. In a separate experiment, we induced endometriosis using wild-type donor endometrium and wild-type recipients, and simultaneously performed adoptive transfer of FACS sorted GFP+ LpM (collected from a naïve mouse) into the cavity of recipient mice. As before, lesions were collected and sorted into GFP+ (lesion-resident LpM) and GFP-monocytes / macrophages. In a separate experiment, we also identified that both Tim4+ and Tim4-LpM trafficked into lesions (Fig.S3). mRNA concentrations of TAM-like and SAM-like markers were evaluated. The TAM-like marker *Folr2* exhibited highest mRNA concentrations in lesion-resident macrophages derived from the endometrium (p<0.05; Fig2B)*. Gas6* mRNA was most abundant in cells derived from infiltrating monocytes*. Mrc1* exhibited higher expression in lesion-resident macrophages derived from the endometrium and infiltrating monocytes (Fig.2B). The SAM-like markers *Spp1* (p<0.05) and *Mmp12* (p<0.05) exhibited increased expression in endometrial-derived macrophages compared to peritoneal derived macrophages. *Ccr2* mRNA concentrations were also elevated in both endometrial-derived lesion-resident macrophages (p<0.01) and the GFP-host-derived population (p<0.05) compared to peritoneal-derived lesion-resident macrophages. Thus, the data are consistent with donor endometrial macrophages and GFP-infiltrating populations being predominantly monocyte-derived as inferred by the scRNA-Seq data presented above (EM and LMono clusters and Cousins et al^27^). Our data indicate that monocyte-derived cells appear to give rise to pro-disease TAM and SAM-like cells. Conversely, the lesion-resident macrophages that are derived from LpM^19^ possess a mature macrophage phenotype and the qPCR data are consistent with these cells having limited differentiation capacity within the lesion. Using immunofluorescence, TAM-like (LRM4; Gas6) and SAM-like (LRM3; Spp1) macrophages were detected *in situ* in mouse (Fig.2C) and human (Fig.2D-E) lesions.

**Figure 2:**
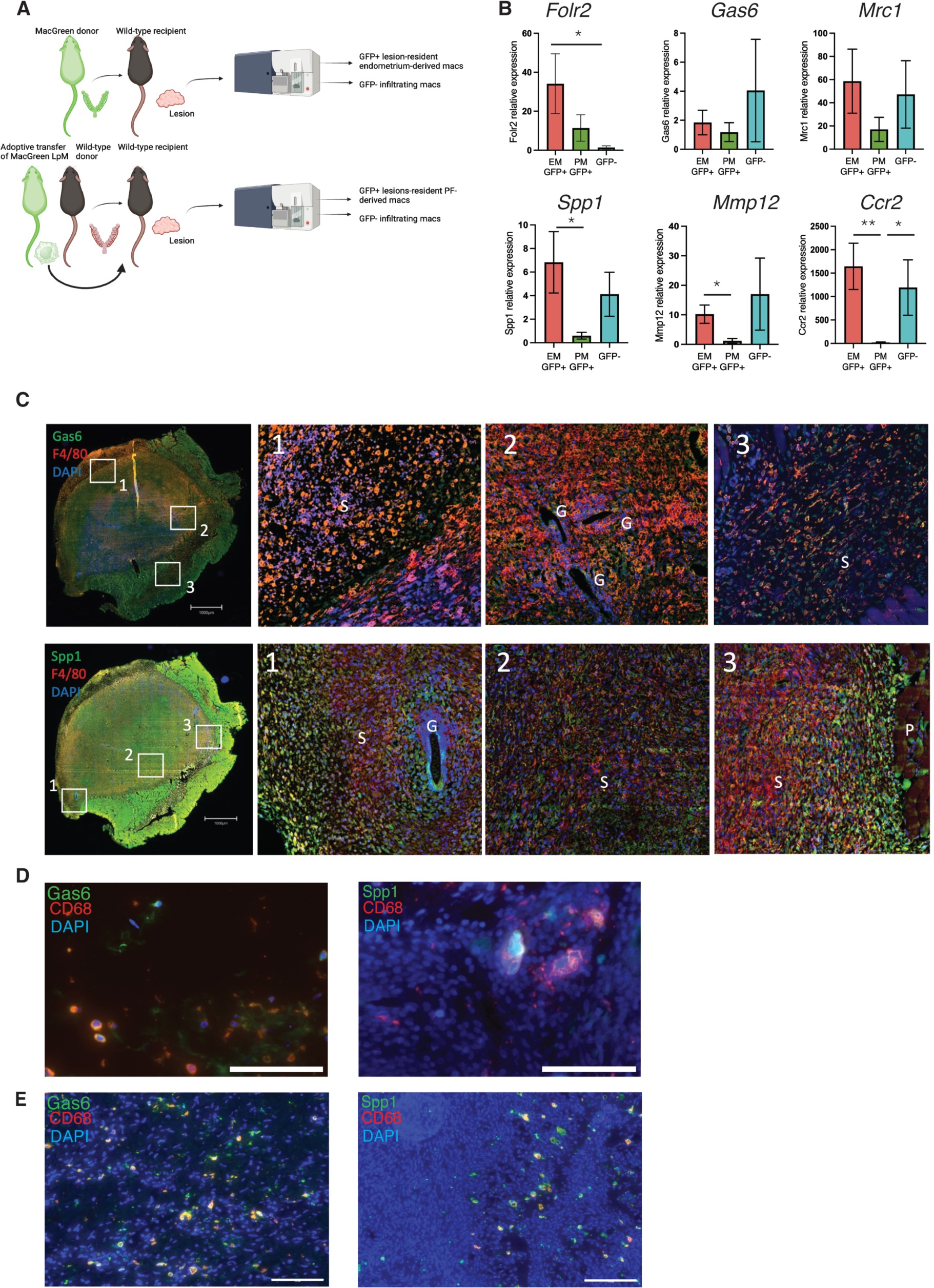
SAM-like and TAM-like cells appear to arise from monocyte precursors and not infiltrating peritoneal macrophages. A) Schematic representation of fate-map experiments that enable FACs isolation of macrophages from different locations. To isolate endometrial-derived macrophages from lesions (n=6 mice), ‘menses’-like endometrium from MacGreen donor mice is transferred into the peritoneal cavity of wild-type recipients and lesions allowed to develop for 14 days. Lesions were recovered and GFP+ endometrial-derived macrophages are isolated. The GFP-macrophage fraction was constituted from monocytes that extravasate from lesion blood vessels and differentiate into macrophages and infiltrating peritoneal macrophages (n =6). To isolate peritoneal macrophages from lesions we performed adoptive transfer of LpM (F4/80+) isolated from the peritoneal lavage of MacGreen mice and injected into the peritoneal cavity of wild-type recipient mice at the same time as endometrium from wild-type mice. Lesions formed over 14 days, followed by FACs isolation of GFP+ (peritoneal) lesion-resident macrophages (n=6). The GFP-fraction was also collected and was constituted by endometrial-derived macrophages and extravasated monocyte-derived macrophages (n=6, collectively n=12 for the GFP-fraction). B) Relative mRNA concentrations of TAM-like (*Folr2*, *Gas6* and *Mrc1)* and SAM-like (*Spp1* and *Mmp12)* macrophages markers assessed by QPCR. C) Immunolocalization of Gas6 and Spp1 (green) and colocalization with F4/80 (red) in mouse lesions (G; gland, S; stroma, P; peritoneum). D-E) Immunolocalization of Gas6 and Spp1 (green) and colocalization with CD68 (red) in lesions recovered from women with endometriosis (D: peritoneal lesions, E: endometrioma). Scale bar = 100μM. Statistical analysis was performed using a Kruskall-Wallis and a Dunn’s multiple comparison test. *: p<0.05, **: p<0.01.

### Lesion-resident Folr2+ macrophages exhibit ‘pro-disease’ properties

To begin to ascertain whether the TAM-like macrophages identified in the scRNA-Seq dataset exhibited ‘pro-disease’ properties in line with their transcriptional phenotype, we FACS sorted Folr2+ macrophages from the peritoneal lavage and lesions of mice with experimentally induced endometriosis (Fig.3A). A greater proportion of lesion-resident macrophages were Folr2+ compared to peritoneal macrophages (Fig.3B). Isolated Folr2+ and Folr2-macrophages were cultured *in vitro* to obtain conditioned media (CM) which was then used in downstream functional assays. Initially, we exposed human endometrial stromal cells to macrophage CM for 3 days followed by qPCR on extracted RNA. Macrophage CM had no impact on the expression of the proliferation marker *Mki67*, whereas CM from lesion-resident Folr2+ macrophages (but not lesion-resident Folr2-or peritoneal macrophages) induced elevated mRNA concentrations of *Col1a1* and *Tgfb1* (p<0.05; Fig.3C). This was consistent with pathophysiological processes taking place within lesions such as extracellular matrix deposition / fibrosis and trans-differentiation. Next, human umbilical vein endothelial cells (HUVECs) were plated on Matrigel® in trans-well plates and exposed to macrophage CM. Multiple parameters were evaluated including branches, segments, junctions and meshes (Fig.3D). We found that CM from lesion-resident Folr2+ macrophages induced a significant and rapid increase in angiogenesis (meshes) at 6h (Fig.3E, p<0.01), whereas CM from lesion-resident Folr2-macrophages did not. Both Folr2+ and Folr2-peritoneal macrophages induced a significant increase in angiogenesis (meshes) at 8h and 6h, respectively.

**Figure 3:**
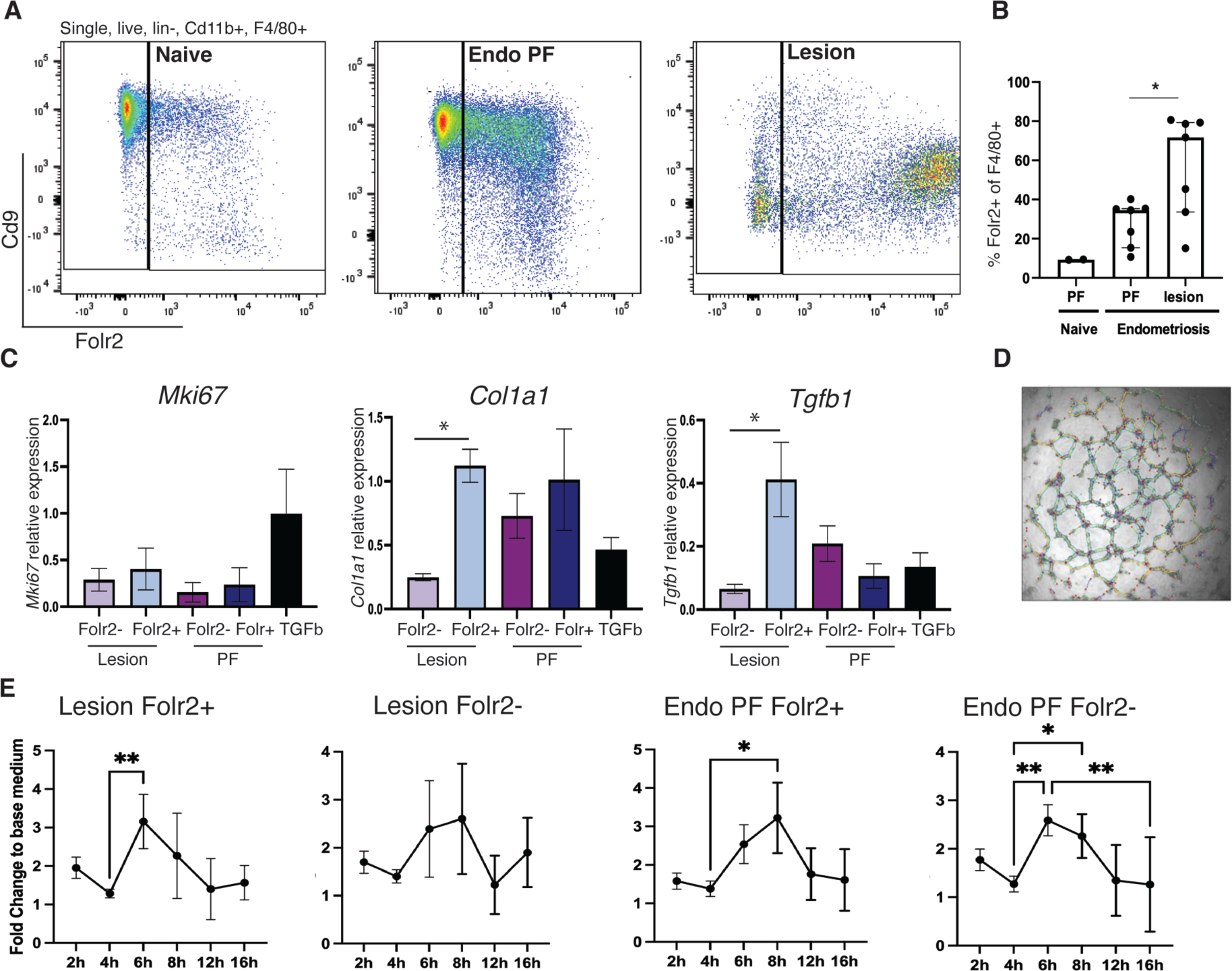
Lesion-resident Folr2+ macrophages exhibit ‘pro-disease’ properties. A) Representative flow plots showing Folr2 expression on F4/80+ macrophages isolated from PF from naïve mice, and PF and lesions recovered from endometriosis mice (n=7 mice). B) Quantification of flow data, showing numbers of Folr2+ macrophages in PF and lesions. FACs sorted Folr+ and Folr2-macrophages derived from PF and lesions of mice with endometriosis were cultured and conditioned media (CM) collected (24h). C) To assess the impact of endometriosis-associated Folr2+ macrophages on proliferation and ECM deposition / fibrosis, primary human (eutopic) endometrial stromal cells (ESCs; from women without endometriosis) were exposed to macrophage CM (24h; diluted 1:1 in minimal media) and the mRNA concentration of *Mki67*, *Col1a1* and *Tgfb1* assessed by QPCR. Human umbilical vein endothelial cells (HUVECs) plated onto Matrigel were also exposed to macrophage CM and network formation assessed. D) Representative image of endothelial cell networks (meshes, branches, junctions, nodes and segments were quantified). E) Quantification of meshes suggests that lesion-resident Folr2+ induce a rapid increase in angiogenesis compared to lesion-resident Folr2-, whereas PF Folr2-macrophages have a more potent effect on angiogenesis compared to PF Folr2+. Statistical analysis was performed using a Kruskall-Wallis and a Dunn’s multiple comparison test. *: p<0.05, **: p<0.01.

### A unique population of monocyte-derived LpM is evident in mice with experimental endometriosis

Our previous work suggested that monocyte-derived LpM confer protection against development of endometriosis lesions. In experiments that limited recruitment of monocytes to the peritoneal cavity, significantly more lesions developed, whereas re-programming the cavity such that embryo-derived LpM were replaced by monocyte-derived LpM resulted in significantly fewer lesions^19^. However, in our scRNA-Seq studies, the altered immune environment induced by surgery (abundant transient LpM in both Endo-Ovx and Sham; Fig.1 and Fig.S2) decreased the distinction with which transcriptional alterations in peritoneal macrophages could be attributed to endometriosis. Thus, we repeated the experiment using ovary-intact mice (Endo-Intact and Naïve). UMAP projection revealed 15 clusters and cell identity was assigned based on expression of canonical markers (Fig.S4). In Endo-Intact mice, an almost entirely unique population of monocyte-derived LpM was evident (Fig.4A and B) and characterised by the presence of *Ccr2* and loss of *Timd4* (Fig.4D). Identification of this population using scRNA-Seq further substantiates our previous findings that endometriosis triggers monocyte recruitment and heightened monocyte input into the LpM pool^19^, and builds on this data to indicate that monocyte-derived LpM are transcriptionally unique. Monocyte-derived LpM were characterized by DEGs *Saa3*, *Apoe*, *Pid1*, *Ltc4s*, *Cebpb* and *Ccl6* (SI Table 3 for gene list). Unique GO terms for this cluster included ‘positive regulation of cellular process’, ‘cellular response to stimuli’ and ‘regulation of metabolic process’. Unique KEGG terms included ‘lipid and atherosclerosis’, ‘sphingolipid signalling pathway’, ‘endocytosis’ and ‘oestrogen signalling pathway’ (SI Table 4). This population shares several DEGs with the transitory LpM population in the Ovx dataset, and *Apoe* is a consistent DEG across transitory LpM, LpM2 (Ovx dataset; Fig.1 and Fig.S2) and monocyte-derived LpM (Intact dataset). LpM1 exhibits a comparable prototypical transcriptional phenotype in the two datasets, whereas LpM3 (Intact dataset) clusters separately from LpM1 but shares many of the same tissue-resident markers.

**Figure 4:**
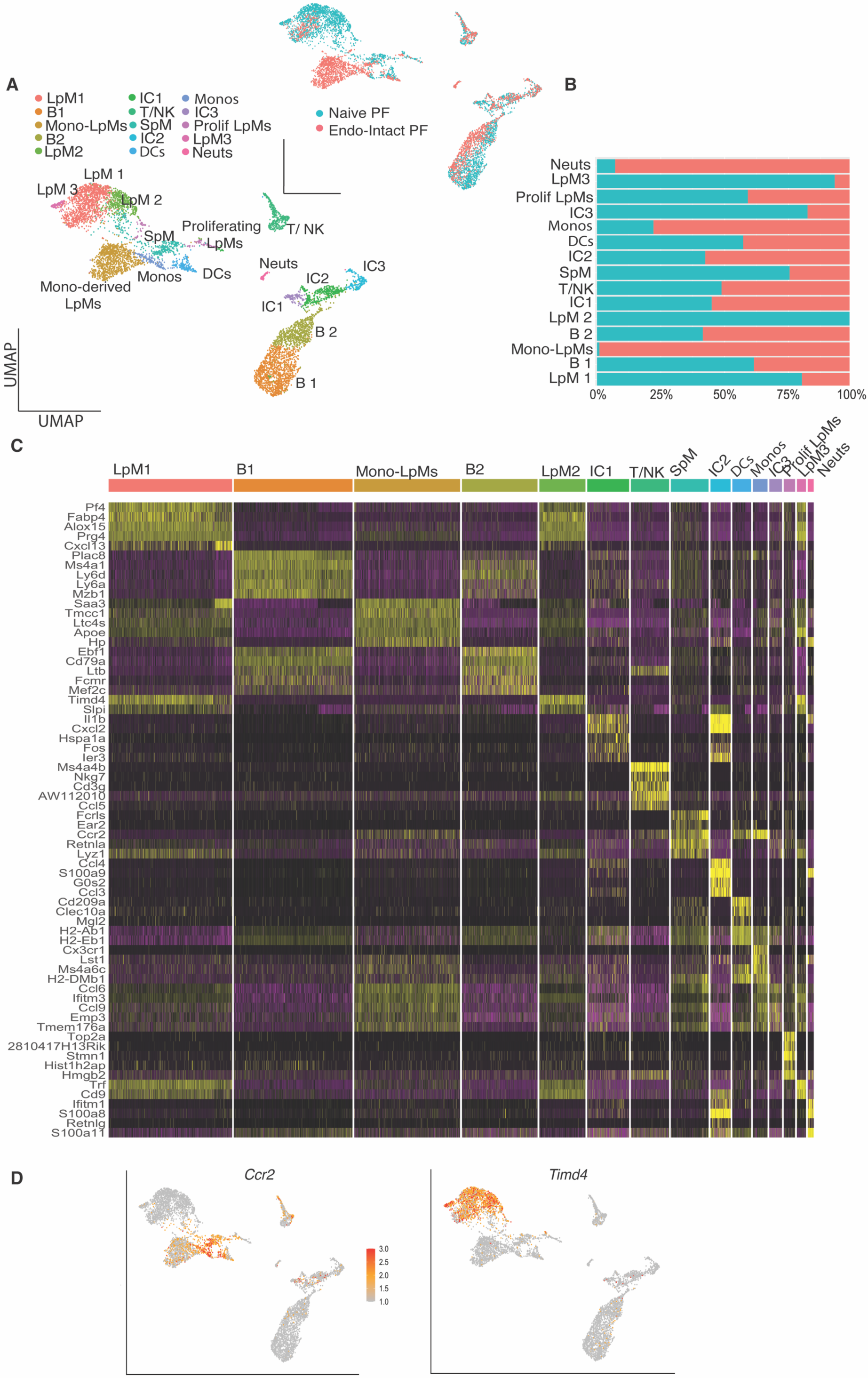
A unique population of monocyte-derived LpMs is evident in mice with experimental endometriosis. A) UMAP projection of CD45+ isolated from the peritoneal lavage from naïve mice (Naïve PF; n=5) and mice with endometriosis (ovaries intact; Endo-Intact PF; n=5). Inset shows UMAP based on library ID. B) Bar chart showing cluster membership of different samples. C) Heatmap showing top 5 DEGs per cluster. D) Feature plots of *Ccr2* and *Timd4* expression.

### Apoe regulates peritoneal macrophage populations and limits growth of lesions in experimental endometriosis

Initially, we presumed that the protective mechanism of monocyte-derived LpM must be a result of increased phagocytic capability. However, in FACS sorted monocyte-derived LpM vs prototypical LpM (Tim4-vs Tim4+) isolated from Endo-Intact and naïve mice, we found no significant differences in phagocytosis (Fig.S5). *Pid1*, *Saa3, Apoe* and *Lrp1* were key DEGs in the monocyte-derived LpM population (Fig.5A), and qPCR analysis also confirmed an up-regulation of these genes in Tim4-peritoneal macrophages, isolated from mice with induced endometriosis compared to those without (Fig.5B). Of the key genes, *Apoe* appears to have an implicit role in LpM function in endometriosis, thus we aimed to perform a gain of function experiment whereby exogenous Apoe (mimetic) was delivered into the peritoneal cavity of mice with experimentally induced endometriosis (intact mice); the Apoe mimetic (COG-133, 3μM in 200μl sterile H2O) or vehicle were delivered daily (via intra-peritoneal injection), with injections initiated at the same time as ectopic tissue transfer and for 2 weeks following. Lesions were monitored using non-invasive bioluminescent imaging (Fig.5C). Treatment with the Apoe mimetic significantly reduced the bioluminescent signal (Fig.5D; p<0.05) and cross-sectional area (p<0.05) of lesions. Flow cytometry revealed a loss of LpM in endometriosis mice exposed to daily i.p injections of vehicle, consistent with the expected macrophage disappearance reaction. Conversely, mice exposed to the Apoe mimetic exhibited significantly more Tim4+ LpM (Fig.5E-F; p<0.05). Numbers of recruited SpM were similar in both Apoe and vehicle treated mice. In pulmonary fibrosis, Apoe produced by monocyte-derived macrophages plays a key role in the resolution of established lesions in the lung. In this context, Apoe was found to promote phagocytosis of type 1 collagen by macrophages in an LRP dependent manner^28^. Thus, we used Masson Trichrome staining to evaluate the amount of collagen deposition in lesions collected from vehicle treated mice with endometriosis, as well as those injected with the Apoe mimetic. Compared to the eutopic endometrium, lesions collected from vehicle treated mice exhibited significantly greater levels of collagen (p<0.01), whereas there was no significant difference in collagen content between eutopic endometrium and lesions from Apoe treated mice (Fig.5G-H).

**Figure 5:**
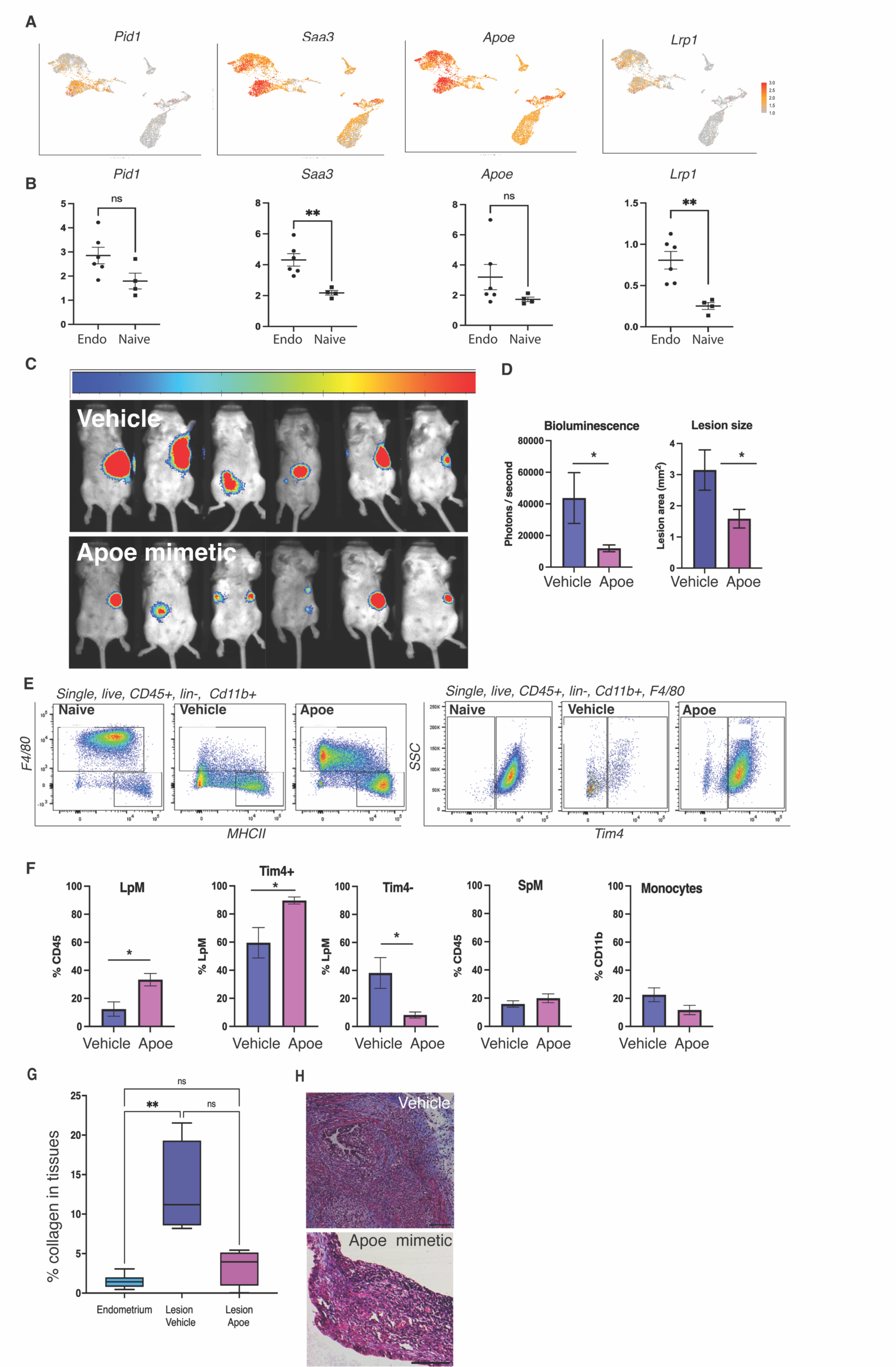
Apoe regulates peritoneal macrophage populations and limits growth of lesions in experimental endometriosis. A) Feature plots of monocyte-derived LpM subpopulation marker genes, *Pid1*, *Saa3*, *Apoe* and *Lrp1.* B) QPCR for marker genes on FACs sorted Tim4-(monocyte-derived) LpM isolated from naïve mice (n=4) and mice with experimental endometriosis (n=6). C) Mice with induced endometriosis were injected i.p with and the Apoe mimetic peptide COG 133 (Generon A1131; 300μM in 200μl dH_2_O, n=6) or vehicle (n=6) daily for 14 days from day of endometriosis induction. Bioluminescent imaging of lesions was performed twice weekly and the images shown are from day 13 of the experiment. D) Quantification of bioluminescent signal on day 13 and quantification of lesion area following histological processing, image capture and measurement using Fiji. E) Representative flow plots of peritoneal lavage showing LpMs (F4/80^hi,^ MHCII^lo^) and SpM (F4/80^lo^, MHCII^hi^; left side) and Tim4+ LpMs (right side) in naïve and endometriosis mice injected with vehicle or Apoe mimetic. F) Quantification of peritoneal macrophages and monocytes (flow data). G) Masson Trichrome stain was performed on lesions collected from endometriosis injected with either vehicle or Apoe mimetic and areas of collagen deposition (blue stain) quantified. H) Representative Masson Trichrome images of lesions. Data (n= 6) are presented as a box plot with the min and max. *: p< 0.05

### Cross-species comparison reveals remarkable concordance in endometriosis-associated macrophage phenotypes and conserved population markers

Design and testing of future macrophage-targeted therapies for endometriosis relies on the understanding of how macrophage subpopulations are similar / unique between the species. Thus, we performed a comparative single-cell transcriptomic analysis of mouse endometriosis-associated macrophages from our preclinical model and their human counterparts. Individual analyses of peritoneal lavage and lesion mouse CD45+ cells from the Endo-Ovx dataset were comparatively analysed with human CD45+ cells from an endometriosis peritoneal fluid dataset (n=1)^29^ and an aggregation of 9 peritoneal endometriotic lesion samples^30^, respectively. After the initial quality control steps and CD45+ filtering, we isolated single-cell transcriptomes from 6026 mouse and 8384 human peritoneal fluid cells. UMAP projection revealed 11 clusters of cells derived from Endo-Ovx mouse peritoneal lavage, with macrophages the predominant immune cell type (Fig.6A-C). Cell populations aligned with our previous aggregate analysis (Fig.1 and Fig.S2) and we detected a new cluster, LpM3, that was not originally identified in this dataset and was characterised by expression of complement receptors and lysozyme 2 (*C1qa, C1qc* and *Lyz2*). In the human dataset, UMAP projection revealed 15 clusters, concordant with the analysis by Zou et al^29^, with similarly predominant macrophages (Fig.6B, C). Initially, MAST analysis was used to identify exact-match DEGs expressed in homologous combined macrophage clusters, with human DEGs converted to mouse orthologs and both filtered by direction of change. Hypergeometric testing revealed a significant overlap between exact-match up-regulated DEGs in the mouse and human peritoneal fluid macrophages, with an 8.5-fold greater enrichment in the overlap than would be expected by chance (hypergeometric p < 0.0001), and a 1.8-fold enrichment in the overlap in down-regulated DEGs (hypergeometric p < 0.005) (Fig.6D). Next, we performed an integrated cross-species analysis of the peritoneal fluid datasets. Cross-species homology mapping was performed and datasets then subset for monocytes / macrophages based on common genes and cross-species integration carried out. UMAP projection of integrated peritoneal fluid datasets composed of 3644 mouse and 5113 human cells revealed 6 populations of monocytes / macrophages (Fig. 6E), with UMAP projection by species showing high concordance between datasets (Fig. S6A). Cluster validation was performed by mapping cell barcodes from specific populations, identified in individual species comparative analysis, onto the integrated UMAP (Fig.S6C-D). Visualisation of clusters (Fig.6E), revealed *Ccr2* to be upregulated on cells that are analogous with mouse SpM (named (S) pM in the integrated dataset) and Vcan+ pM, indicating recent differentiation from monocytes, with highest expression on (S) pM alongside the characteristic high *H2-Aa* and low *Adgre1* expression seen in mouse SpM. *Ccr2* was expressed at a lower level in Vcan+ pM, alongside expression of *S100a8* and *S100a9* (see SI Table 5 for gene list). *Timd4* was up-regulated on prototypical pM and Lyve1+ pM, indicative of long residence. Prototypical pM expressed high *Adgre1* and genes characteristic of steady-state prototypical LpM in mice: *Cxcl13, Prg4, Alox15 and Fabp4.* Lyve1+ pM also expressed scavenger receptors *Marco*, *Cd163,* and complement *C2*. Transitory (Transit) pM also expressed *Lyve1*, although at a lower level, alongside downregulation of *H2-Aa* and *Ccr2*. As previously shown in Fig.1A, a proliferative population (Prolif pM) was identified (*Top2a, Cdk1,* and *Mki67).* Evaluation of cluster occupancy by species revealed proliferative and transitory populations to be consistent, however Lyve1+ pM and prototypical pM populations were expanded in human and mice, respectively (Fig.6F). Of all the clusters, the Lyve+ pM population was significantly under-represented in the mouse dataset suggesting this population is largely unique to the human. Unique GO terms for this cluster included ‘neurogenesis’, ‘neuron projection development’ and ‘neuron development’; others included ‘negative regulation of cellular metabolic process’, ‘phosphorylation’ and ‘regulation of intracellular signal transduction’. Unique KEGG terms included ‘endocytosis’, ‘pathways in cancer’ and ‘lipid and atherosclerosis’ and ‘growth hormone synthesis and secretion’ amongst others (SI Table 6).

**Figure 6:**
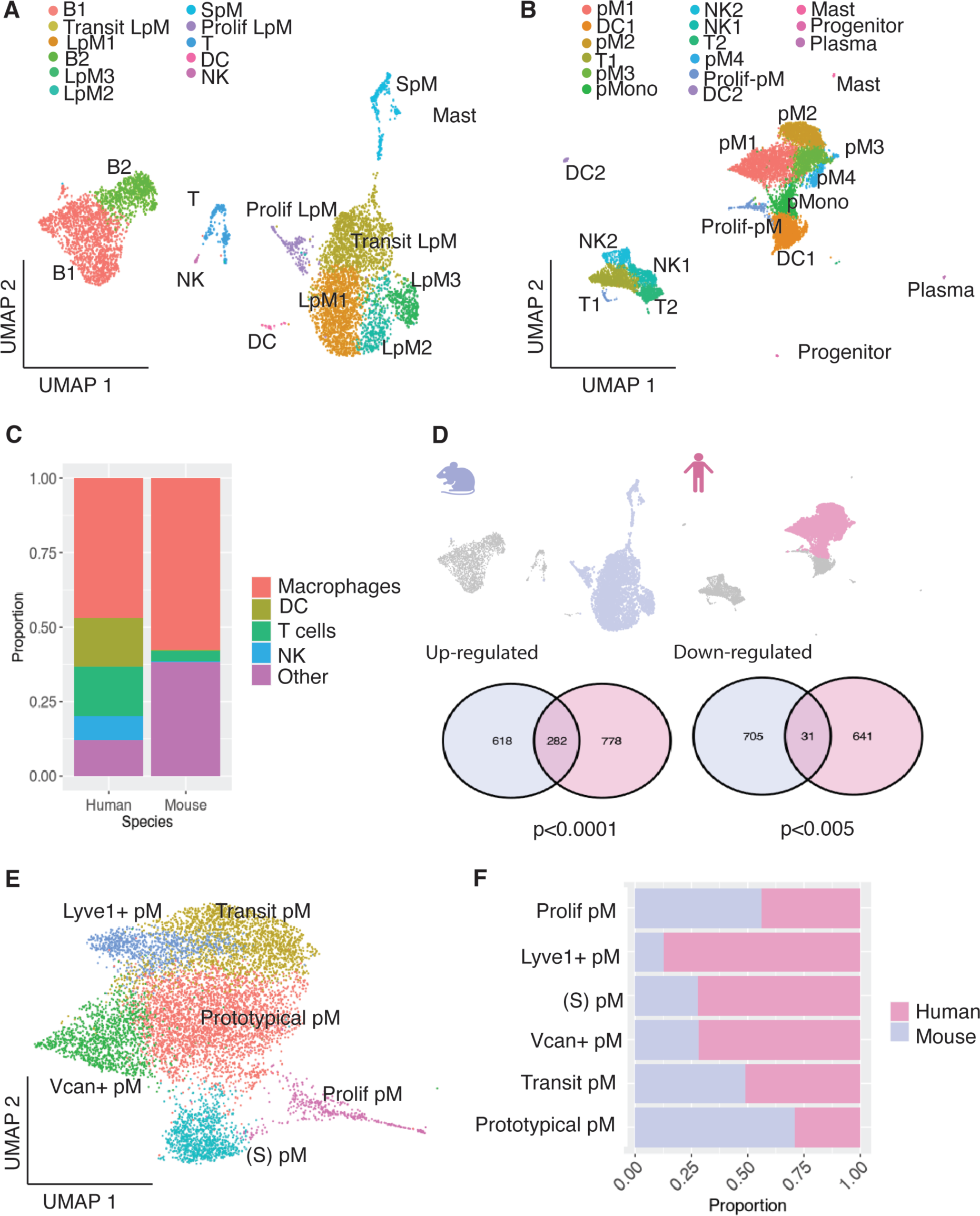
Cross-species mapping of mouse and human peritoneal macrophages. A) UMAP projection of CD45+ cell derived from mouse peritoneal fluid (Seurat v.5), B) UMAP projection of CD45+ cells derived from human peritoneal fluid (Zou et al publicly available dataset). C) Bar chart showing the proportions of each cell type present in the two datasets. The ‘other’ population includes other cells excluding macrophages, DC, T and NK (e.g B cells, mast cells). D) The macrophage subset was extracted from each dataset (lilac and pink for mouse and human, respectively) and evaluated for shared up- and down-regulated genes, see Venn diagrams. E) Cross species integration of single-cell RNA-sequencing data was performed to map mouse and human macrophage subpopulations. F) Bar chart showing cluster member proportions for each species.

In the lesion datasets, we isolated the single-cell transcriptomes from 5494 mouse, and 11965 human CD45+ cells. UMAP projection revealed 13 clusters in the Endo-Ovx mouse lesions, as per our previous aggregate analysis, with macrophages accounting for >75% of cells (Fig.7A-C). In the human endometriosis lesions, 20 clusters were identified, with macrophages far less prevalent and accounting for just over 25% of cells, concordant with prior analysis by Tan et al (Fig.7B-C)^30^. Comparisons of DEGs from the homologous mouse and human lesion macrophage clusters revealed an 11-fold enrichment in the overlap of common up-regulated exact-match DEGs (hypergeometric p < 0.0001), but no significant enrichment of exact-match down-regulated DEGs (Fig.7D). Integrated cross-species analyses of the lesion transcriptomic datasets were performed using one-to-one orthologs as described above. UMAP projection of integrated lesion datasets composed of 4858 mouse and 5517 human macrophages revealed 7 clusters (Fig.7E). Compared to peritoneal macrophages, less alignment was seen between lesion macrophages across the two species (Fig.S7A). Cluster validation was performed as before (Fig.S7B). *Ccr2* was found to be upregulated on a substantial lesion-resident monocyte (Lmono) population, alongside high *H2-Aa* and *Clec4b1* (SI Table 7), indicating these cells were recently monocyte derived. Il1b+ LM expressed high levels of *Il1b,* and several down-regulated genes, suggesting a transitory population. Lyve1+ LM expressed *Lyve1, Folr2, Mrc1* and *Gas6*, indicating a TAM-like gene signature as seen in the mouse dataset. Spp1+ LM expressed *Spp1, Lgals, Trem2,* and *Cd9,* concordant with a SAM-like gene signature. *Lgals* was also expressed by Vcan+ LM, alongside *S100a8/9* expression. Peritoneal LM highly expressed prototypical LpM-like genes, including *Prg4*, *Alox15,* and *Fabp4,* suggesting infiltration of peritoneal macrophages into lesions as previously seen in the mouse and in the human data^30^. Prolif LM expressed many markers of proliferation including *Top2a* and *Mki67.* Evaluation of cluster occupancy by species revealed the SAM-like Spp1+ pM and monocytic populations were relatively consistent, however Vcan+ pM and Lyve1+ pM populations were expanded in humans and mice, respectively (Fig.7F). Further analysis of Vcan+ macrophages derived from lesions are warranted as these appear to be specific to the human. However, GO analysis only returned 3 unique terms: ‘tyrosine phosphorylation of STAT protein’, ‘regulation of tyrosine phosphorylation of STAT protein’ and ‘post transcriptional gene silencing’ and no unique KEGG terms (SI Table 9). These comparative and integrated cross-species analyses highlight the phenotypic concordance between specific endometriosis-associated macrophage populations in humans and a mouse model of experimental endometriosis, while elucidating species-specific population enhancements. Taken together, these results are vital for further studies on the development of macrophage-directed therapies for the treatment of endometriosis.

**Figure 7:**
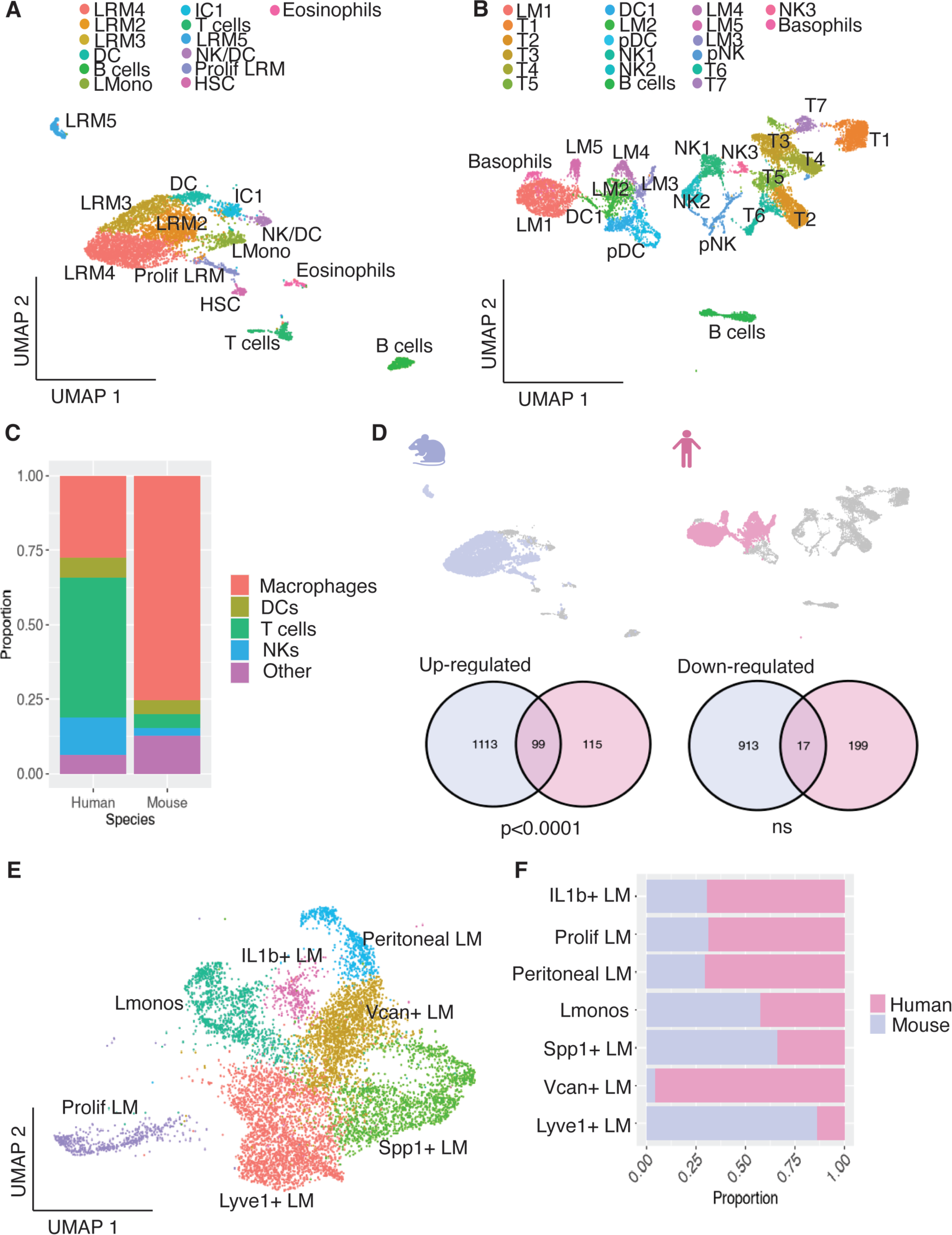
Cross-species mapping of mouse and human lesions-resident macrophages. A) UMAP projection of CD45+ cells derived from lesions recovered from a mouse model of experimental endometriosis (Seurat v.5), B) UMAP projection of CD45+ cells derived from human endometriosis lesions (Tan et al, publicly available dataset). C) Bar chart showing the proportions of each cell type present in the two datasets. The ‘other’ population includes other cells excluding macrophages, DC, T and NK (e.g eosinophils, basophils, B cells). D) The macrophage subset was extracted from each dataset (lilac and pink for mouse and human, respectively) and evaluated for shared up- and down-regulated genes, see Venn diagrams. E) Cross species integration of single-cell RNA-sequencing data was performed to map mouse and human macrophage subpopulations. F) Bar chart showing cluster member proportions for each species.

## Discussion

In the current study, we performed single-cell profiling on macrophages recovered from the endometriotic niche (lesions and peritoneal cavity) of a preclinical model, revealing the full complexity of macrophage phenotype and aligning transcriptomic signatures to macrophages with pro-disease and pro-resolving properties. Here, we shed new light on population-specific markers and reveal the concordance between human and mouse endometriosis-associated macrophages.

scRNA-Seq analysis identified 7 distinct lesion-derived monocyte / macrophage clusters in the model, highlighting heterogeneity of macrophage subpopulations far beyond those previously identified^19^. Some lesion-derived cells clustered with either endometrial or peritoneal macrophages, highlighting that previously identified origins can also be found at the transcriptional level. Of note, a recognisable pattern of monocyte-macrophage transition traditionally associated with wound healing can be observed in the data^31^. In lesions, this cycle is characterised by a significant monocyte population likely recruited from the circulation, an early pro-inflammatory macrophage population (LRM2) and pro-repair phenotypes associated with ECM remodelling (LRM3; SAM-like macrophages) and promotion of cell proliferation and angiogenesis (LRM4; TAM-like macrophages). It is likely that this cycle of recruitment and differentiation within lesions is continuous. Both endometriosis lesions and tumours have been referred to as ‘wounds’ that do not resolve, given their propensity for characteristic inflammation, cell proliferation (regeneration), vascularisation and innervation, and transdifferentiation. Tumour macrophages also exhibit transcriptomic and functional heterogeneity analogous to what we observed in endometriosis lesions and a framework has been established in order to classify the spectrum of TAM phenotypes^32^. Although some key markers (*Spp1*, *Folr2*, *Trem2)* were shared between TAM^32^ and LRM sub-populations, we also observed some dichotomy so the nomenclature was less useful in this study.

Of the three identified ontogenies of lesion-resident macrophages^19^, we have demonstrated that endometrial and recruited monocyte-derived macrophages appear to give rise to pro-disease macrophages, whilst peritoneal-derived macrophages exhibited restricted differentiation capacity in lesions. Recruitment / trafficking of peritoneal macrophages during injury has been consistently observed in different visceral organs. During intestinal injury, F4/80^hi^, GATA6+ LpM accumulate at damaged sites in a Ccr2 independent manner. Once at the site of injury, peritoneal macrophages disassemble necrotic cells and contribute to revascularisation and collagen deposition^33^. As it was noted that the lamina propria of the intestine exhibited an accumulation of Ccr2+ monocytes, whereas peritoneal macrophages infiltrated the muscularis of the intestine which has a lower vascular density, the authors proposed this illustrated the importance of blood flow independent macrophage recruitment for rapid tissue repair. Consistent with this, it may be presumed that peritoneal macrophages play a critical role in establishment of endometriosis lesions, prior to development of neovessels. Whilst LpM trafficking to lesions do not appear to adopt a pro-disease phenotype, GO analysis of prototypical LpM suggests roles in ‘cell-cell interaction’ and ‘migration’, ‘platelet activation’ and ‘sprouting angiogenesis’ consistent with a role in wound repair and aligned with a key role for prototypical LpM in repair of damaged peritoneal lining^34, 35^. Thus, we suggest that LpM are vital for the first step of attaching refluxed endometrial tissue to the peritoneal lining, establishment of lesions and development of a blood supply.

Our previous findings indicated that endometrial macrophages are pro-disease^19^. In this study, our scRNA-Seq data indicate that endometrial-derived ‘macrophages’ are largely constituted of monocytes and early, actively proliferating macrophages. We also demonstrated that monocytes recruited directly to lesions eventually yield a pro-disease phenotype, indicating, for the first time that it is largely lesion-resident monocyte-derived macrophages that promote the growth of lesions, neuroangiogenesis, fibrosis, and ultimate maintenance of the lesion, after the initial LpM-mediated attachment phase. In cancer, the previous dogma that monocyte-derived macrophages recruited directly to the tumour exclusively gave rise to TAMs has been challenged and it is now accepted that in some cancers, embryonic-derived tissue-resident macrophages are major contributors to the TAM pool^36^. In general, high TAM infiltration correlates with poor outcomes given their widely accepted role in promoting angiogenesis, tumour growth, ECM re-modelling, and inhibition of anti-tumour responses, e.g. t-cell-mediated cytotoxicity^37, 38, 39^. However, in some cancers TAM can be associated with enhanced anti-tumour properties^40^. Several studies have now suggested that ontogeny plays a key role in pro-vs anti-tumour activity. In endometriosis, monocyte-derived macrophages appear to adopt two different pro-disease phenotypes which we refer to as TAM-like or SAM-like. *Gas6* was highly abundant in the TAM-like population. By binding to its receptors (Axl, Mertk, Tyro3), Gas6 is known to significantly affect cell cycle progression in cancer cells, promote angiogenesis and modulate the immune environment of the tumour^41^. In other systems this pathway has been intimately linked with the process of fibrosis, including in lung and liver fibrosis^42^. In patients with intra-uterine adhesions (IUA), CD301+ endometrial macrophages exhibited increased abundance and secreted Gas6 that promoted endometrial fibrosis^43^. Ccl8 was also a top DEG in the TAM-like population and has previously been shown to promote progression of tumour cells, induce invasion and stem-like traits in glioblastoma^44^, and is also negatively correlated with patient prognosis in a several different cancers^45^. It is also known to be induced by lactate in TAM^46^, consistent with altered metabolism and increased lactate levels in both cancer^47^ and endometriosis^48^. TAM-like macrophages also exhibited restricted and elevated expression of Folr2, a reliable marker induced in both lesion-resident TAM-like macrophages and some peritoneal macrophages of mice with endometriosis. Folr2 expression is associated with pro-tumour macrophages that exhibit an anti-inflammatory and immunosuppressive microenvironment in several cancers ^49^, however in some indications (e.g. breast cancer) has been associated with CD8+ T cell infiltration and better patient survival^49^. In this study, isolated lesion Folr2+ macrophages exhibited a pro-angiogenic and pro-fibrotic role, consistent with expression of other ‘pro-disease’ genes in this population. The second pro-disease population identified in lesions exhibited a scar-associated macrophage (SAM-like) phenotype, analogous to macrophages that control fibrosis in the liver^50, 51^, lung^52^, heart, kidney and skin^22^ and this phenotype appears to be conserved across species and tissues and exhibits specificity to fibrotic disease^22, 52^. The SAM-like population is characterised by expression of *Spp1, Gpnmb, Cd63* and *Trem2,* and *Mmps*, and is predominantly monocyte-derived in other tissues with a role in extracellular matrix deposition and remodelling^51^.

Of note, whilst we have identified that monocyte-derived macrophages give rise to lesion-resident macrophages with pro-disease phenotypes, our previous studies using Ccr2^-/-^ mice demonstrated that loss of this recruitment axis resulted in development of more and larger lesions^19^ as a result of a depleted monocyte-derived LpM pool. This also highlights distinct differences between lesion-resident and peritoneal cavity-resident macrophages in endometriosis. Akin to cancer, we have identified that cellular ontogeny accounts only for some of the transcriptional heterogeneity, thus signals received from the local environment are also major determinants of phenotype^53, 54^.

Peritoneal cavity macrophages exhibit a highly specific transcriptional profile that is distinct from most lesion-resident macrophages. Consistent with previous scRNA-seq studies investigating the transcriptome of peritoneal macrophages in steady state^6^, we observed 3 LpM populations corresponding to the most abundant prototypical LpM as well as differentiation intermediates in our naïve dataset. Conversely, in our Endo-Intact dataset a monocyte-derived LpM population was identified, congruent with the ‘protective’ population of monocyte-derived LpM that we previous identified in our preclinical model and induced in endometriosis^19^. Characteristic DEGs include *Pid1, Saa3, Apoe,* and *Lrp1*, each of which are associated with glucose and lipid metabolism and cholesterol efflux and may also play a role in fibrosis. Interestingly, studies have suggested that total cholesterol and triglycerides are upregulated in the serum of women with endometriosis^55^ and are positively correlated with disease severity^56^. As cholesterol is a key precursor of steroid hormones, this points to a link between the increased oestradiol biosynthesis in endometriosis lesions^57^ and cholesterol metabolism in the peritoneal cavity. Thus, we postulate that the protective monocyte-derived LpM population reduces availability of triglyceride and cholesterol availability for lesion growth, as well as subsequently lowering local oestrogen biosynthesis, thus reducing drivers for endometriosis maintenance and progression. Future work will focus on evaluating this hypothesis. In support of our findings, Jenkins et al reported that inflammatory macrophages abundant during mild zymosan induced inflammation expressed genes that overlapped with LpM of recent monocyte origin in female mice including Apoe^7^.

In this study, we have used two variations of experimental endometriosis: Endo-Ovx (our original model) and Endo-intact. We included the intact model following the realisation that the transcriptomic profile of monocyte-derived LpM was masked in the Ovx model due to the large population on transitory LpM (monocyte-derived) present in both Sham and Endo-Ovx mice. These findings are consistent with previous studies demonstrating that surgery alters the immune composition and peritoneal macrophage dynamics in mice^6^ and we suggest that future studies take this into consideration in their experimental design. *Apoe* was a robust marker of monocyte-derived LpM in both variations of the endometriosis model. In a gain-of-function experiment utilising an Apoe mimetic peptide we found that the size of endometriosis lesions was reduced. We identified reduced collagen in Apoe treated lesions in support of previous studies indicating that Apoe can promote phagocytosis of type 1 collagen by macrophages in an LRP dependent manner^28, 58^. Aside from its role in lipid metabolism, Apoe is also known to modulate both adaptive and innate immune responses in mice^59^, and as the Apoe mimetic significantly elevated numbers of Tim4+ LpM it may be suggested that Apoe has an autocrine role in regulating differentiation of Tim4-LpM into Tim+ LpM in the peritoneal cavity. Significant questions remain regarding its role in lipid metabolism and cholesterol efflux in the peritoneal cavity and the pathogenesis of endometriosis.

A recent scRNA-Seq study on human endometriosis-associated macrophages by Tan et al, identified heterogenous subpopulations of macrophages^30^, and key macrophage populations identified in our mouse model are also present in the human endometriosis ecosystem. The concordance between human and mouse was remarkable, with a significant number of shared markers, indicating that we have a clinically tractable model for our study. For example, *FOLR2* and *MRC1* were top markers in their LYVE+ lesion-resident macrophages, indicating similarities with our TAM-like macrophages. *SPP1* was also a top marker in the activated macrophage population, and this population aligns with our SAM-like population. Similar to our data, Tan et al also identified infiltrated macrophages (with high expression of *CCR2)* and peritoneal macrophages (*ICAM2* and *FN1*). To further validate the concordance of mouse and human lesion-resident macrophages we performed integrative cross-species mapping^60^. The analysis demonstrated co-clustering of the major subpopulations for both human and mouse and highlighted proportions in each cluster. Overall, the similarity was beyond what we had predicted, but some deviations were apparent. For example, *SPP1+* (SAM-like) macrophages were expanded in the human, whereas *LYVE*+ (TAM-like) macrophages were expanded in mice. *VCAN*+ macrophages appeared to be almost uniquely human, and future work should focus on determining the function of these cells. In the peritoneal fluid, we have highlighted key similarities in macrophage populations between human^29^ and mouse, with the identification of equivalent populations to mouse LpM (prototypical tissue resident population), transitory (differentiation intermediates), and monocyte-derived SpM. In the human, a population of *LYVE*+ peritoneal macrophages are present but significantly under-represented in the mouse data, indicating these may play a more substantial role in human peritoneal physiology and future work should characterise function.

In summary, we have demonstrated that macrophage sub-populations are transcriptionally diverse, and that the major pro-disease macrophage populations in endometriosis lesions are congruent with TAM and SAM and are largely monocyte-derived. We have identified that peritoneal macrophages are distinct from lesion-resident macrophages and exhibit both prototypical and protective phenotypes, with macrophage-derived Apoe as a key mediator of protection against development of endometriosis lesions. In the future, we propose that endometriosis researchers should look to cancer and other fibrotic diseases to exploit and repurpose macrophage-directed therapies that target these specific phenotypes and simultaneously modify the immune environment to promote accumulation of pro-resolving macrophages.

## Acknowledgements

The authors thank Prof. Neil Henderson, Beth Henderson and Dr Prakash Ramachandran (University of Edinburgh; Centre for Inflammation Research) for access to the 10X Chromium Controller, help and advice with sample processing and some initial interpretation. We also thank the QMRI flow cytometry and cell sorting facility technicians (University of Edinburgh) for advice on panel design, and cell sorting. We would also like to thank Dr Pavle Vrljicak for critical training and advice on performing bioinformatics analysis and interpretation (University of Warwick). This work was supported by a Medical Research Council (MRC) Career Development Award (MR/M009238/1; to E.G), an MRC Project Grant (MR/S002456/2; E.G), an MRC Centre for Reproductive Health studentship to C.H, a WMS studentship to Y.H, an MRC-DTP in Interdisciplinary Biomedical Research studentship to N.M.A and an MRC-DTP iCASE studentship to I.M. The authors declare no conflict of interest.

## Author contributions

E.G conceived experiments, performed experiments, analysed data, and wrote manuscript and acquired funding; K.P and Y.H conceived and performed experiments, analysed data, and wrote manuscript; I.M performed cross-species analysis and wrote manuscript. P.D performed experiments; C.H conceived and performed experiments; A.C and A.S performed experiments and analysed data; N.M.A performed experiments and analysed data; M.R performed experiments and analysed data. S.O provided critical training and advice on bioinformatics analysis and interpretation and co-supervised Y.H. A.W.H and E.T.C provided feedback on experimental design and manuscript preparation. A.W.H co-supervised C.H.

## Materials and Methods

### Animals and reagents

Wild-type C57BL/6JOIaHsd and FVB female mice were purchased from Harlan (Harlan Sprague Dawley Inc, Bicester, UK) at 8-12 weeks of age. B6.Cg-Tg(Csf1r-EGFP)1Hume/J (MacGreen) express enhanced green fluorescent protein (EGFP) under control of the Csf-1r promoter^61^ and FVB-Tg(CAG-luc,-GFP)L2G85Chco/J were bred in house^62^. Mice were maintained at the University of Edinburgh or the University of Warwick. All animal work was licensed and carried out in accordance with the UK Home Office Animal Experimentation (Scientific Procedures) Act 1986 and the work licensed under PPL 70/8731 and PP0568394 (E.G). Mice had access to food and water ad libitum and were kept at an ambient temperature and humidity of 21°C and 50% respectively. Light was provided 12 hours a day from 7am-7pm. To visualize bioluminescent endometriosis lesions, the substrate D-luciferin (1.5 mg/100 μl in PBS; Sigma-Aldrich, Dorset, UK) was injected s.c. prior to imaging prior to imaging on under anaesthesia using a PhotonIMAGER (Biospace Lab, Paris, France)^62^. For Apoe experiments, mice were injected intra-peritoneally with COG133 (Generon A1131; 3μM in 200μl sterile H20) or vehicle (sterile H20) daily from onset of endometriosis to 14 days post induction.

### Mouse model of induced endometriosis

Endometriosis was induced in mice using a syngeneic model as previous described^16^. The model aims to mirror the process of ‘retrograde menstruation’. In brief, donor mice were induced to undergo a ‘menses’-like event by removing the ovaries and exposing the mice to a hormonal schedule similar to a truncated menstrual cycle and a stimulus that causes the endometrial stromal cells to undergo decidualization^63^. Following P4 withdrawal the endometrial lining begins to shed. 4-6hrs after withdrawal of P4, the ‘menses’-like endometrium was collected and injected into ovariectomized mice supplemented with oestradiol valerate^63^. Lesions were recovered that contain stoma +/-epithelial cells and immune cell influx^16^. For some experiments endometrial tissue or FACS sorted LpM were isolated from MacGreen mice and transferred to wild-type recipients during induction of endometriosis to allow the isolation of endometrial-derived or peritoneal-derived macrophages to be isolated from resultant lesions as previously described^19^. For some experiments recipient mice were left with their ovaries intact and did not receive exogenous oestradiol valerate. For non-invasive bioluminescent imaging of lesions, endometrial tissue from CAG-luc-eGFP mice was injected into the peritoneal cavity of wild-type FVB mice. Lesions were collected 2 weeks following tissue injection into ice-cold DMEM. Peritoneal lavage was recovered by injecting 7 ml ice-cold DMEM into the peritoneal cavity followed by gentle massage and recovery.

### Flow cytometry

Lesions were dissected, pooled from each mouse and placed in 2ml ice-cold DMEM. Tissues were cut into small pieces using a scalpel and digested with 1 unit of Liberase DL, 1 unit of Liberase TL (Roche) and 0.6 mg DNAse enzymes. The tissue and enzymes were incubated for 45 minutes at 37°C, with vortexing every 5 minutes. Following digestion, samples were filtered through 100μM filters. Red blood cells were lysed from peritoneal lavages and cells derived from the endometrium or lesions, and approx. 10^6^ cells per sample were blocked with 0.025mg anti-CD16/32 (clone 93; BioLegend, San Diego, CA, USA) and then stained with a panel of antibodies shown in SI Table 9. Brilliant^TM^ violet stain buffer (BD Biosciences) was included when required. Fluorescence minus one (FMO) and unstained controls were used to validate gating strategies. Just prior to analysis on the flow cytometer, DAPI and 123count eBeads (Thermo Fisher Scientific) were added to samples. Samples were analysed using an LSRFortessa with FACSDiva software or FACSMelody with Chorus software (BD Biosciences) and analysed with FlowJo v.9 software (FlowJo, Ashland, OR, USA). Analysis was performed on single, live cells determined using scatter height vs. area and negativity for live or dead (DAPI or alternative viability dye). For fluorescent activated cell sorting red blood cell lysis, Fc blocking and fluorescent staining were performed as previously described and samples sorted into pure cell populations based on cell surface marker expression using a FACS Aria Fusion (BD Biosciences).

### Single cell RNA-Sequencing and analysis

From donor endometrium (+4-6 hours P4 withdrawal; n=5 mice), endometriosis lesions (lesions from n=10 mice) and peritoneal lavage from Sham (n=5) and Endometriosis mice (n=5), 200,000 CD45+ cells were FACS sorted into 1ml PBS + 2% FBS using a FACS Fusion. Cells were barcoded using a 10X Genomics Chromium Controller^TM^ using established pipelines. Libraries were sequenced by Edinburgh Genomics using a NovaSeq 6000 sequencing system (Illumina®, San Diego, USA). Initial processing was performed using Cellranger (v2.1.1) mkfastq and count (aligned to mouse assembly mm10). For each dataset (filtered data from Cell Ranger 27 pipeline), potentially low-quality cells were filtered out using dataset-specific thresholds. For the Endo-Ovx dataset cell ranger metrics were as follows: estimated number of cells: 1,306, 5,645, 6,720, 6,006 (Menses-like endometrium, Sham PF, Endo-Ovx PF and Lesions, respectively); mean reads per cell: 393,957, 97,631, 77,378, 81,492, median genes per cell: 2,050, 1,326, 1,702, 1,160, sequencing saturation: 93.7%, 90.6%, 84.4%, 92%. For the Endo-Intact dataset, cell ranger metrics were: estimated number of cells: 4,612, 3,069 (Naïve PF and Endo-Intact PF, respectively); mean reads per cell: 148,623, 235,681; median reads per cell: 1,304, 1,133; sequencing saturation: 92.8%, 94.8%. Clustering and analysis of differential gene expression (DGE) was performed using Seurat (v4.4.0 – v5.0.0) in R (v4.3.2). Preferentially, we selected cells where at least 200 genes were detected and only genes that were expressed in at least 3 cells were included in down-stream analysis. To reduce the risk of including low-quality cells we integrated our datasets and performed quality control to filter out cells with high mitochondrial gene expression (8%), followed by data normalisation and scaling as per standard Seurat workflow (Stuart and Satija, 2019). As the samples in each dataset were all processed on the same day, cartridge and sequenced on the same lane, batch correction was not utilised. To visualize the population dynamics and reduce dimensionality of our merged dataset, we ran a principal component analysis (PCA) on the normalized gene matrix of the top 4000 most variable genes. Cell clustering was visualized using Uniform Manifold Approximation and Projection (UMAP; PCAs 11, dims 11, resolution 0.6 (aggregate, Fig.1), PCAs; 17, dims; 17, resolution 0.6 (PF; Fig.3)). The function Clustree^64^ was used to visualise the relationships between clusters and how these changed at different resolutions. This also allowed for the identification of distinct and unstable clusters. The Seurat function ‘FindAllMarkers’ was used to identify marker genes for each cluster in the UMAP projection according to the inbuilt Wilcox statistical test. To identify potential doublets from our dataset, we used the ‘doublet finder’ function to exclude these cells and repeated normalisation, feature selection, scaling and UMAP projection and cluster marker identification as described above. Ultimately, we retained 1295 cells from donor endometrium, 5560 cells from lesions, 4966 cells from sham peritoneal fluid and 6265 from endometriosis peritoneal fluid (Endo-Ovx). For the intact dataset we retained 3895 (Naïve PF) and 2583 (Endo-Intact).

### KEGG and GO

The statistical analysis and visualisation of functional profiles for each gene cluster was analysed using the function compareCluster from the package *clusterProfiler*. First, data sets were subset to include only the macrophage subpopulations. Differentially expressed genes (DEGs) were then generated using Seurat’s FindAllMarkers function and subset for markers that were upregulated with adjusted p values lower than 0.05. This list was used for the geneCluster argument of compareCluster. For the identification of specific Gene Ontology (GO) and Kyoto Encyclopedia of Genes (KEGG) terms, a custom universe was also created which included only genes that were expressed in the subset data and this list was used in the universe argument of compareCluster. For GO enrichment analysis and KEGG enrichment analysis, the fun argument was set to enrichGO or enrichKEGG respectively.

### Cross-species analysis

Analyses were performed on mouse lesion and peritoneal lavage data from the Endo-Ovx dataset above, and human data retrieved from the publicly available datasets of Tan et al (peritoneal endometriotic lesions)^30^ and Zou et al. (endometriosis peritoneal fluid)^29^ using the guideline best practices for single-cell analysis described by Heumos et al^65^. Raw scRNA-seq data was demultiplexed, aligned to reference genomes (h38, mm10) and processed using Cell Ranger Software as above. Analysis of gene-barcode matrices were performed in Seurat (5.0.0) in R (4.3.2) with Seurat objects (v5) created using cells with > 200 genes, and genes expressed in > 3 cells. Quality filtering of scRNA-seq data was performed at the per sample level to remove cells of low quality using multiple filtering parameters of mitochondrial percentage (< 20), number of genes detected (> 350), gene expression counts (> 1000 in human samples; > 350 in mouse samples due to reduced sequencing depth), and log10GenesPerUMI (> 0.8). The packages scDblFinder, SoupX, and cc.genes (Seurat) were used to identify and remove doublets, estimate ambient RNA contamination, and perform cell cycling scoring, respectively^66, 67^.

#### Parallel analysis

Seurat package was used to normalise individual expression matrices using the ‘NormalizeData’ and ‘ScaleData’ functions, with the ‘FindVariableFeatures’ function implemented to select the top 2000 variable genes for PCA analysis within the ‘RunPCA’ function. The 9 human (peritoneal) lesion Seurat objects were integrated to batch correct for inherent patient differences, the other 3 datasets are composed of individual Seurat objects. The Seurat functions ‘RunUMAP’, ‘FindNeighbours’ and ‘FindClusters’ were used for visualisation and clustering. The parameters for clustering lesions were as follows: mouse; 30 PCAs, 1:30 dims, resolution 0.5, human; 30 PCAs, 1:30 dims, resolution 0.65; combined; 50 PCAs, 1:30 dims, resolution 0.4. The parameters for PF were as follows: mouse; 30 PCAs, 1:30 dims, resolution 0.4, human; 30 PCAs, 1:30 dims, resolution 0.7, combined; 50 PCAs, 1:30 dims, resolution 0.3. As the mouse peritoneal fluid and lesion datasets contained only CD45+ cells, human datasets were subset on CD45+ expressing clusters, followed by re-running of visualisation and clustering steps. Seurat ‘FindAllMarkers’ Wilcoxon rank-sum tests implemented in a one-versus-all fashion were used to identify marker genes for manual annotation according to canonical biomarkers identified in literature study. DEG analysis was performed using the MAST functionality in Seurat ‘FindAllMarkers’ to account for within-sample correlation. Hypergeometric tests to compute the significance of overlapping DEGs between mouse and human datasets were performed in R (4.3.2) on homologous DEGs with a > 1.5 or < -1.5 log2 fold change, against the hypergeometric distribution conferred by random iterative sampling of the overlap occurring when sampling the same number of DEGs from each species-specific homologous genomic background in 100000 simulations to account for the different number of genes original R objects.

#### Integrative cross-species mapping

Cross species integration of single-cell RNA-sequencing data was performed in Seurat 5.0.0 using the Seurat V4 CCA O2O strategy described by Song et al. due to the high integrated scoring of species mixing and biological conservation^60^. Homology mapping was performed using one-to-one orthologous genes between human and mouse samples. Prior to creation of Seurat objects, human genes within the gene-barcode matrices were translated to their mouse one-to-one orthologs using the convert_human_to_mouse_symbols function in the nichenetr R package^68^. Human genes without mouse one-to-one orthologs were removed, and all datasets were then subset on genes common to both human and mouse datasets. Seurat objects were then created, processed, and visualised as prior, with integration of human and mouse macrophage clusters performed using Seurat v4 anchor based CCA integration, followed by marker gene and DEG analyses as described above. For validation of integrated clustering, the Seurat ‘WhichCells’ functionality was used to identify cell barcodes of specific clusters in the single-species analyses, with these cells then projected onto UMAPs in the cross-species integrated analyses.

### Human samples

Endometriotic lesion biopsies (peritoneal and endometriomas) were collected from patients enrolled in the EndoWar study (University of Warwick REC: 19/LO/1647) who provided informed consent and were undergoing laparoscopy for suspected endometriosis. Tissues and clinical metadata were collected in accordance with the WERF EPHect guidelines^69, 70^.

### Cell culture

#### Generation of conditioned media from FACS sorted macrophages

Folr2+ and Folr2– macrophages were isolated by fluorescent-associated cell sorting (FACS) from peritoneal fluid and lesions from mice with experimental endometriosis (n=7) using a BD FACS Fusion Aria cell sorter. Sorted macrophages were seeded at a density of 10,000 cells per 50 µL of recovery media (DMEM-hi-glucose, 2% HI-FBS, 25 mM HEPES, 50 µM β-mercaptoethanol, 1mM Sodium Pyruvate, 1x antibiotic-antimycotic, 1x non-essential amino acids), and cultured for 24 hrs at 37°C (5% CO2) humid incubation. Following overnight macrophage cell recovery culture, the media was aspirated, and the macrophages were replenished with fresh low-serum media (DMEM-hi-glucose, 0.2% HI-FBS, 25 mM HEPES, 50 µM β-mercaptoethanol, 1mM Sodium Pyruvate, 1x antibiotic-antimycotic, 1x non-essential amino acids) to create conditioned medium. Macrophage conditioned media was then harvested after 24 hrs overnight incubated culture at 37°C (5% CO2) humidity and cryopreserved at –80°C for later use.

#### HUVEC cell culture

Human umbilical vein endothelial cells or HUVECs were derived from a pool of donors (Promocell, #C-12203) and expanded to passage 5 in VEGF-rich complete media (Promocell, #C-22011) at 37°C, 5% CO2, humidified incubation.

#### Endometrial stromal cell culture

Human endometrial stromal cells (huESCs) were derived from two participants, as previously described^81^. Endometrial biopsies were received from Jan Brosens (University of Warwick). Accordingly, different endometrial cellular components were dissociated by mincing the tissue with a scalpel blade for 5 minutes then digested by shaking incubation in an enzymatic cocktail consisting of 5mL phenol red-free Dulbecco’s Modified Eagle Medium (DMEM)/F12, 0.5 mg/mL collagenase I and 0.1 mg/mL DNAse I (Sigma) at 37°C, 5% CO2, humidity for 1 hour. Human ESCs were then specifically separated from the resulting homogenate by washing in complete growth medium (DMEM/F12 containing 10% dextran-coated charcoal stripped FBS (DCC-FBS), 1% penicillin-streptomycin, 2 mM L-glutamine, 1 nM E2 (Sigma-Aldrich) and 2 mg/ml insulin (Sigma-Aldrich) followed by filtration through a 40 µm cell strainer, where huESCs were obtained in the flowthrough. The flowthrough was further washed in complete growth medium and centrifuged (400xg 5 minutes, RT). Resulting cell pellets were resuspended in 10 mL complete growth medium and seeded in tissue culture flasks. Any contaminating (non-adherent) cell were removed by replacing the medium following overnight cell culture. The cell culture medium was subsequently replenished at 48-hour intervals and sub-confluent monolayers of huESC were passaged at a 1:3 ratio using 0.25% Trypsin-EDTA.

### Ex vivo functional assays

#### Phagocytosis assay

Mouse peritoneal lavage was collected and prepared for FACS with blocking and staining steps as previously described. Samples were stained with a panel of antibodies as show in SI Table 9. The cell suspension (100μl) at a concentration of 1-5x10^6^cells/ml was incubated with a 1:100 final dilution of latex beads-rabbit IgG-FITC complex from the Cayman Chemical Phagocytosis Assay Kit (IgG FITC) for 1 hour at 37°C in FACS tubes. To assess phagocytic activity, the suspensions were pelleted at 400 x g for 5 minutes and resuspended in 300μl of flow buffer (PBS+2%BSA) before being analysed on the LSR Fortessa™ with FACS Diva software and analysed with FlowJo as previously described. The gating strategy live, single, CD45+, lineage-, Cd11b+, MHCII^lo^, F4/80^hi^ was used to identify the large peritoneal macrophages (LpM). Monocyte-derived LpM were Timd4^lo^ and embryo-derived LpM were Timd4^hi^ and sorted using this distinction. The internalisation of the latex beads was quantified as a measure of macrophage phagocytosis by noting the fluorescence intensity of FITC and compared between monocyte and embryo-derived (long-lived) macrophages.

#### Endometrial stromal cell gene expression

A collation of 20,000 huESCs derived from two participants were seeded in complete growth medium into a 96-well plate and allowed to expand for 48 hours at 37°C, 5% CO2, humidified incubation. This was followed by overnight serum starvation in low serum growth medium (DMEM/F12 containing 2% dextran-coated charcoal stripped FBS (DCC-FBS), 1% penicillin-streptomycin, 2 mM L-glutamine, 1 nM E2 (Sigma-Aldrich) and 2 mg/ml insulin (Sigma-Aldrich)). Following overnight serum-starvation, huESCs were cultured in the presence of macrophage conditioned medium (CM) derived from macrophages isolated from mice with endometriosis and naïve controls over a predetermined period of 3 days. Macrophage CM was pre-diluted at a ratio of 1:4 in low-serum huESC culture medium. Untreated and TGF-β1 (5 ng/mL, 10 ng/mL)-treated cells were used as negative and positive controls, respectively, and were cultured in stock CM culture medium diluted 1:4 in low-serum huESC culture medium. HuESCs were cultured in macrophage conditioned media for 3 days at 37°C, 5% CO2, humidified incubation before harvesting.

#### Angiogenesis assay

HUVECs underwent VEGF-starvation for 24 hrs prior to use in an angiogenesis assay (Promocell, #C-22010). A fluorescent Calcein stain in Ca2+ and MG2+ free HBSS (40 minutes, 37°C, 5% CO2 humidified incubation) was applied to HUVECs before seeding 15,000 cells resuspended in low-serum media (0.5% FBS and VEGF-negative, Promocell #C-22010 diluted 1:3 in basal media # C-22010B) on to a layer of Matrigel BME (25 µL, Corning, #356231) in a 96-well transwell culture arrangement. Conditioned media (200 µL) was diluted 1:9 in low-serum (0.5% FBS and VEGF-negative) media, and added to the lower chamber. The formation of vascular networks were imaged at timepoints 2, 4, 6, 8, 12 and 16 hrs on an EVOS M7000 Imaging system. Phase contrast images were then analysed in Fiji using the pre-validated Angiogenesis Analyzer workflow, as previously described^82^.

### Real-time qPCR

RNA was extracted from FACs sorted macrophages using RLT(lysis) buffer and an RNAeasy Kit (Qiagen, Hilden, Germany). Concentration and purity were assessed using a Nanodrop 1000 (Thermo Fisher Scientific). For FACS sorted cells RNA was amplified cDNA synthesized using a SeqPlex RNA Amplification kit (Merck Life Science, UK) as per manufacturer’s instructions. For endometrial stromal cells, cDNA was synthesized using SuperScript Vilo Enzyme (Thermo Fisher Scientific) with 100 ng starting template in a 20μl reaction. A standard curve was generated by pooling samples and performing four 10-fold dilutions. PCRs (10μl) were performed using validated Taqman^TM^ qPCR assays (20mM; Applied Biosystems, UK) and Express qPCR Supermix (Thermo Fisher Scientific). cDNA was added at 1μl per reaction and thermal cycling conditions were performed on a 7900 Fast real-time PCR machine (ThermoFisher Scientific) in 384-well plates with technical duplicates performed. 18S (Thermo Fisher Scientific) was selected as the reference gene. Data were analysed using the relative standard curve method, and samples were normalized to 1 consistent sample.

### Immunofluorescence

Immunofluorescence was carried out as previously described^19, 21, 71^. In brief, 5 µm thick sections of FFPE lesions collected from the mouse Endo-Ovx endometriosis model were stained by dual immunofluorescence to identify F4/80+GAS6+ TAM, and F4/80+SPP1+ SAM. Sections were adhered to microscopic slides by overnight incubation at 65°C. After dewaxing and rehydrating in gradient solutions of ethanol, sections underwent antigen retrieval in trypsin (Sigma, #, 1g/ml, 10 minutes, 37°C). Sections were permeabilised in 0.25% triton for 30 minutes at room temperature (RT). Endogenous peroxidases were blocked in 3% hydrogen peroxidase (30 minutes, RT), and non-specific binding was completed using normal goat serum (NGS) (30 minutes, RT). The first primary antibody (rat anti-F4/80 (eBioscience, #14-4801-82, dilution 1:100)) was applied overnight, followed by F4/80 antigen detection using a goat polyclonal anti-rat antibody conjugated to horseradish peroxidase (IMPRESS, # MP-7404), followed by a 10-minute incubation in Tyramide red (1:50 in diluent). Sections underwent a repeat antigen retrieval that was optimised for the second primary antibody in the dual immunofluorescence strategy. Gas6 antigens were retrieved in citrate buffer (pH6). Spp1 antigens were retrieved in Tris-EDTA buffer (pH8). As before, sections were blocked for endogenous peroxidases and non-specific binding before applying primary antibodies rabbit polyclonal anti-Gas6 (ThermoFisher, # PA5-79300, dilution 1:100) and rabbit polyclonal anti-Spp1 (ThermoFisher, # PA5-34579, dilution 1:100). Gas6 and Spp1 antigen detection was achieved using a goat polyclonal anti-rabbit antibody conjugated to horseradish peroxidase (IMPRESS, # MP-7451), followed by a 10-minute incubation in Tyramide green (1:50 in diluent). Washes were completed in PBS-Tween in between steps. Sections were finally mounted in an anti-fading mounting medium with DAPI (Vectashield, # H-2000). Antibodies were validated using single stain controls, and the omission-of-one primary antibody controls in a dual stain. Images of whole lesions were captured using a 20X objective on an EVOS M7000 (Invitrogen, UK) imaging system.

Dual immunofluorescence to identify CD68+GAS6+ TAM and CD68+SPP1+ SAM in human lesions was performed as follows: Sections (3µm) were deparaffinised and antigen retrieval performed in TRIS-EDTA buffer pH9. After endogenous peroxide blocking (using 3% hydrogen peroxide) and species-specific blocking (using normal goat serum 5% in 2% bovine serum albumin), sections were stained with monoclonal mouse anti-human CD68 clone PG-M1 - 1:100 (Dako Omnis: GA61361-2). Second antigen retrieval and blocking were performed as described above and subsequent staining was performed with either rabbit anti-Gas6 - 1:100 (PA5-79300) or rabbit anti-SPP1 - 1:100 (PA5-34579). Secondary antibody and fluorescent staining was performed using the ImmPRESS IgG polymer detection kits (MP-7451, MP-7452) and Tyramide TSA plus kit with either Fluoresciene or Cy5. Autofluorescence quenching was performed with the TrueBlack® Lipofuscin Autofluorescence Quencher (Biotium: 203007) according to manufacturer’s instructions. Slides were mounted with VECTASHIELD PLUS Antifade Mounting Medium with DAPI (Vector Laboratories: H-1000-10) and imaged on an EVOS M7000 (Invitrogen, UK) microscope.

### Masson-trichrome stain

Paraffin-embedded mouse lesion and endometrium tissue sections (3 µm) were stained using the Abcam Trichrome kit (ab150686 Trichrome stain) following the protocol provided.

### Fiji analysis

Masson-trichrome images were obtained using the Leica DMi8 with an 10X objective. Collagen intensity analysis was ascertained in Fiji image analysis software (Image J v1.53) by isolating stained area of blue colouration (collagen) using colour thresholding, creating a mask of this selection and utilising particle analysis to measure the total area of the mask. This was then presented as a proportion of the total area of the lesion.

### Statistical analysis

Statistical analysis was carried out in GraphPad Prism 10.0. Data were analyzed for normality using an Shapiro-Wilk normality test. If data were normally distributed, either an ANOVA with a Tukey’s post-hoc test (more than two samples) or a t test (two samples) was performed. If data were not normally distributed, nonparametric tests were used, either Kruskal–Wallis with a Dunn’s post hoc test (more than two samples) or a Mann– Whitney U test (two samples). Statistical significance was reported at p<0.05.

### Data Availability

The sequencing data will be deposited in the GEO repository (https://www.ncbi.nlm.nih.gov/geo/) under accession number (TBC upon publication). Code will be available on Github upon publication.

## Notes

### Competing Interest Statement

The authors have declared no competing interest.

