## Supplementary Figures for "Single-cell analysis identifies distinct macrophage phenotypes associated with pro-disease and pro-resolving functions in the endometriotic niche"

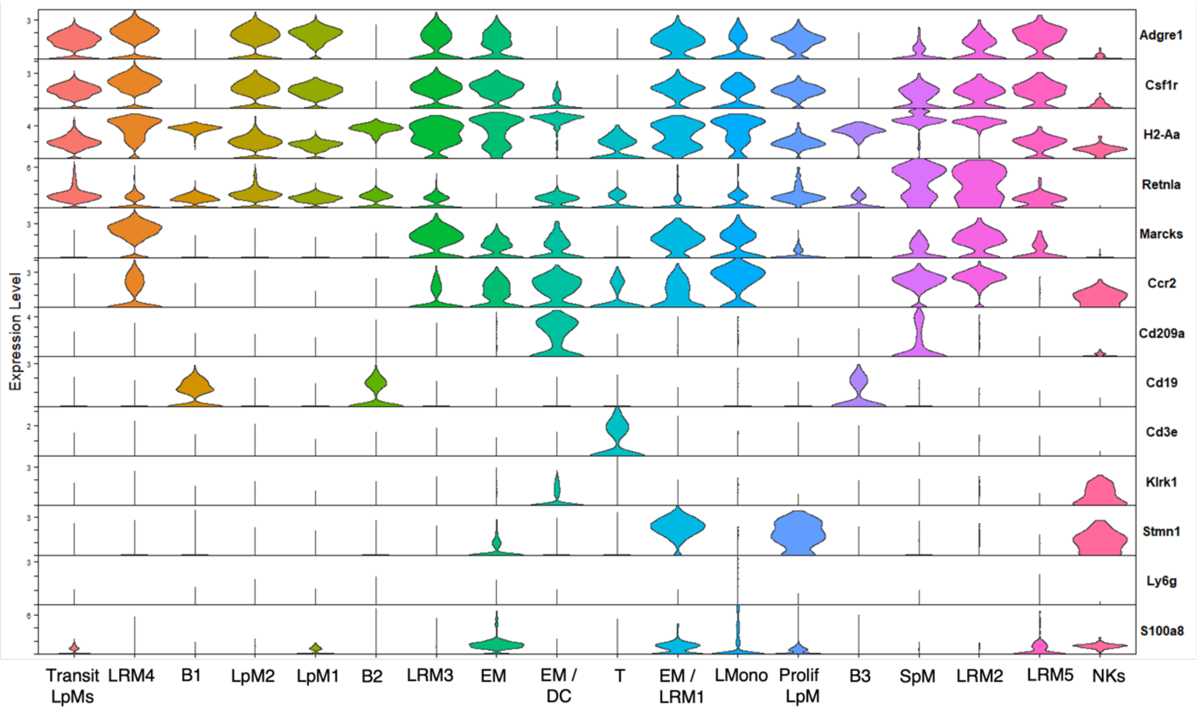


**Figure S1: Canonical markers used for assigning cell identity.** Violin plots showing expression of canonical markers used to identify cell clusters in Figure 1.

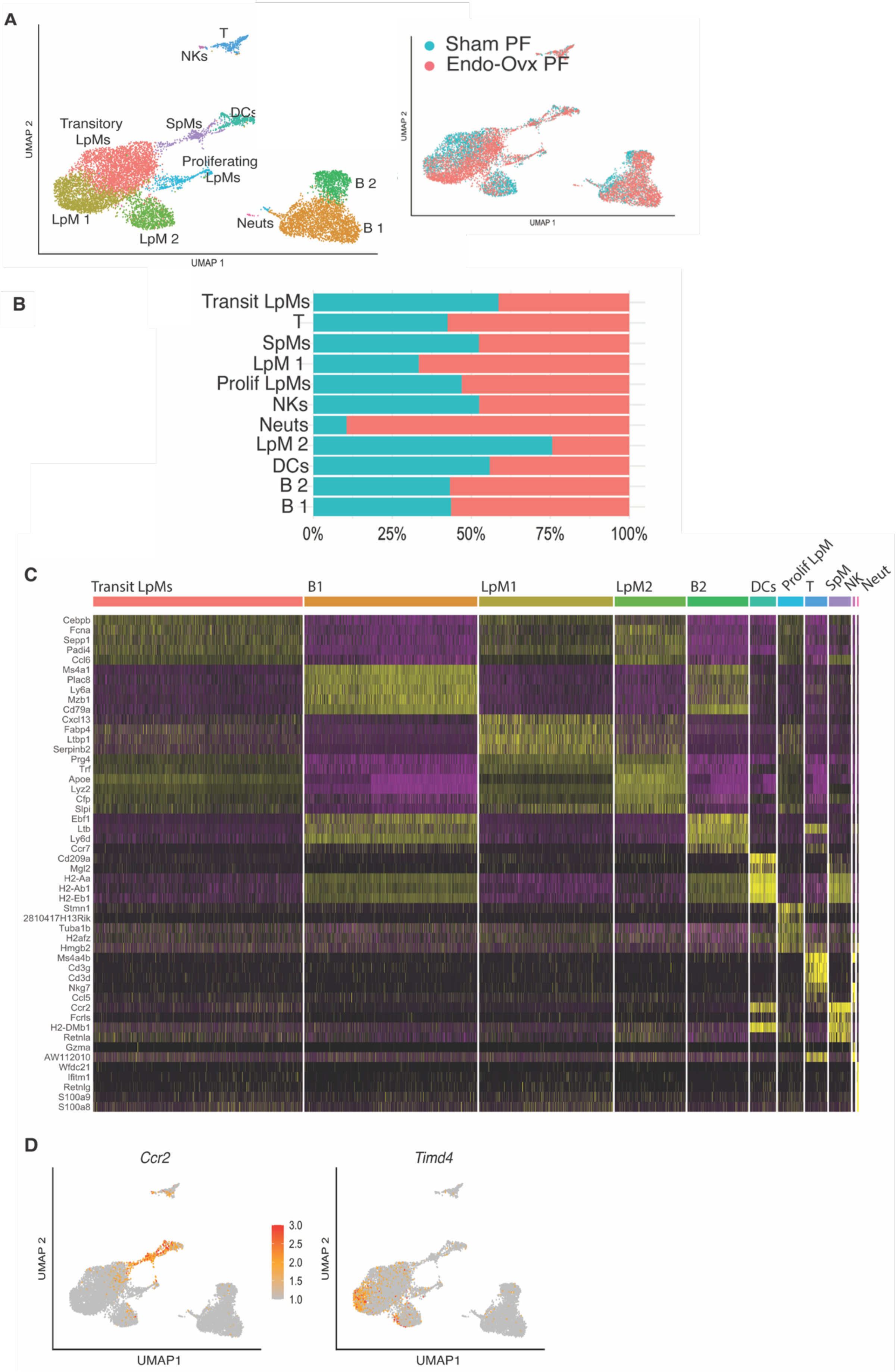
A
T
NKs
Transitory
LpMs
SpMs
DCs
Proliferating
LpMs
B 2
B 1
Neuts
LpM 1
LpM 2
UMAP 2
UMAP 1
Sham PF
Endo-Ovx PF
UMAP 2
UMAP 1
B
Transit LpMs
T
SpMs
LpM 1
Prolif LpMs
NKs
Neuts
LpM 2
DCs
B 2
B 1
0%
25%
50%
75%
100%
C
Transit LpMs
B1
LpM1
LpM2
B2
DCs
Prolif LpM
T
SpM
NK
Neut
Cebpb
Fcna
Sepp1
Padi4
Ccl6
Ms4a1
Plac8
Ly6a
Mzb1
Cd79a
Cxcl13
Fabp4
Ltbp1
Serpinb2
Prg4
Trf
Apoe
Lyz2
Cfp
Slpi
Ebf1
Ltb
Ly6d
Ccr7
Cd209a
Mgl2
H2-Aa
H2-Ab1
H2-Eb1
Stmn1
2810417H13Rik
Tuba1b
H2afz
Hmgb2
Ms4a4b
Cd3g
Cd3d
Nkg7
Ccl5
Ccr2
Fcrls
H2-DMb1
Retnla
Gzma
AW112010
Wfdc21
Ifitm1
Retnlg
S100a9
S100a8
D
Ccr2
Timd4
UMAP 2
UMAP1
3.0
2.5
2.0
1.5
1.0
UMAP 2
UMAP1

**Figure S2: Transcriptomic heterogeneity of peritoneal macrophages in an ovariectomised model of induced endometriosis.** A) UMAP projection of CD45+ cells isolated from peritoneal lavage recovered from Sham and Endo-Ovx mice. Inset shows UMAP based on library ID. B) Bar chart showing cluster membership of different sample types. C) Heatmap showing top 5 DEGs for each cluster. D) Feature plot of *Ccr2* and *Timd4*.

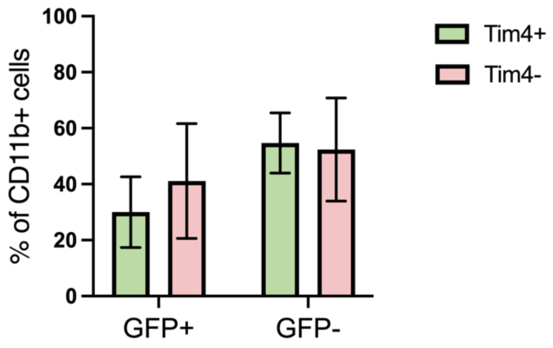


**Figure S3: Adoptive transfer of Tim4+ and Tim4- LpMs reveals that both population sare incorporated into endometriosis lesions.** Tim4+ or Tim4- LpM recovered from MacGreen mice were adoptively transferred into wild-type mice inoculated with wild-type endometrium and GFP+ and GFP- cells recovered from lesions after two weeks. Graph shows quanitification of cells recovered (Tim4+ n=7 mice, Tim4- n=3 mice).

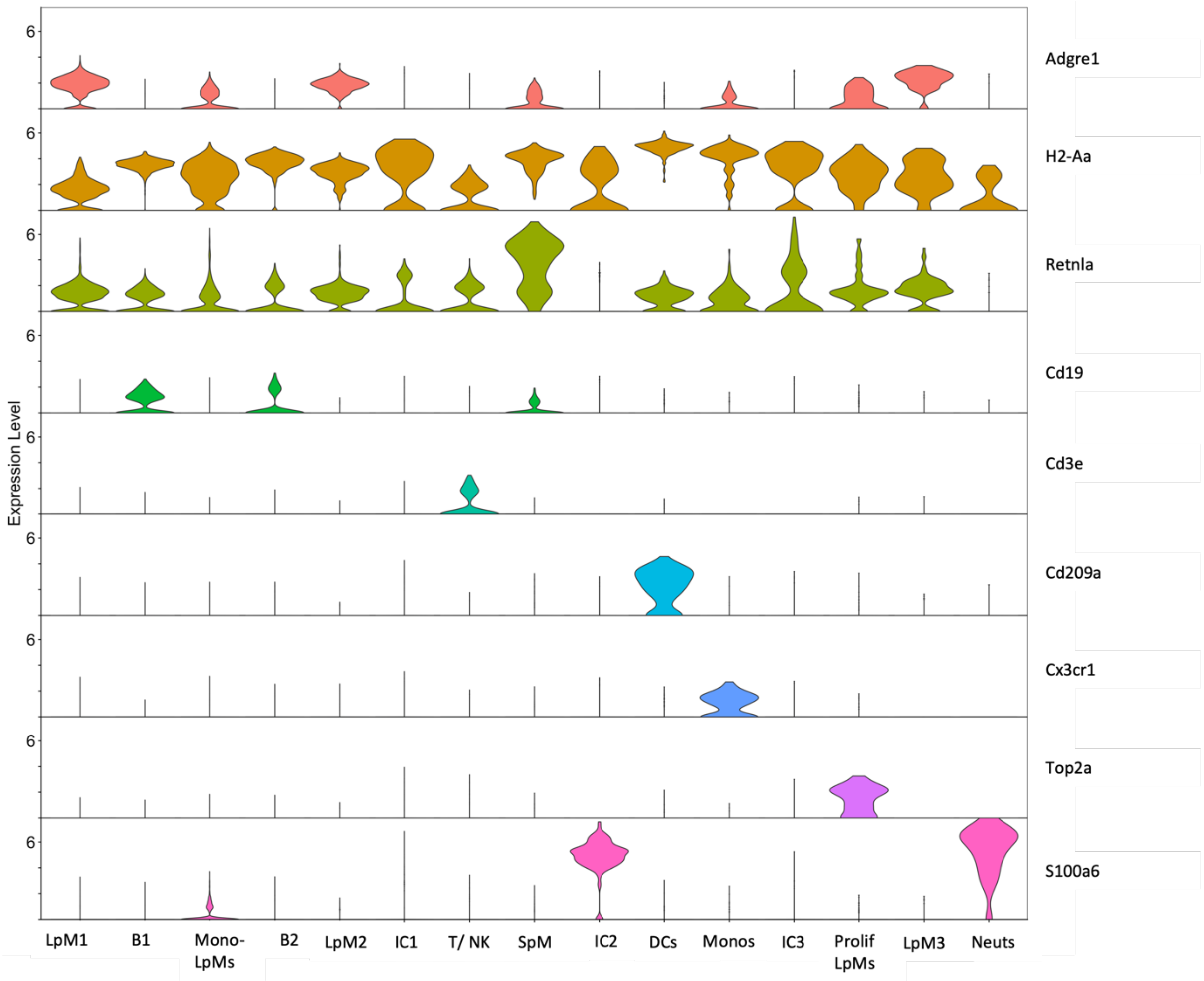


**Figure S4: Canonical markers used for assigning cell identity.** Violin plots showing expression of canonical markers used to identify cell clusters in Figure 3.

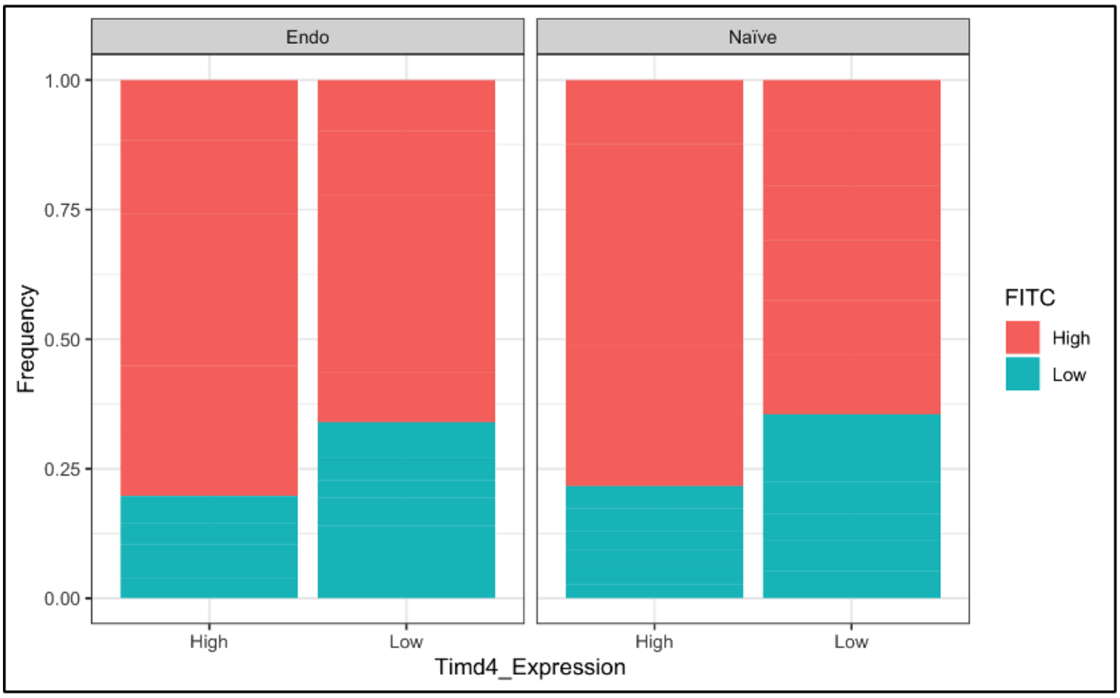


**Figure S5: Analysis of phagocytic activity in Tim4$^{hi}$ and Tim4$^{lo}$ macrophages isolated from the peritoneal cavity of mice with and without induced endometriosis.** Tim4$^{hi}$ and Tim4$^{lo}$ peritoneal macrophages were FACs sorted and incubated with FITC beads and phagocytic uptake measures using flow cytometry.

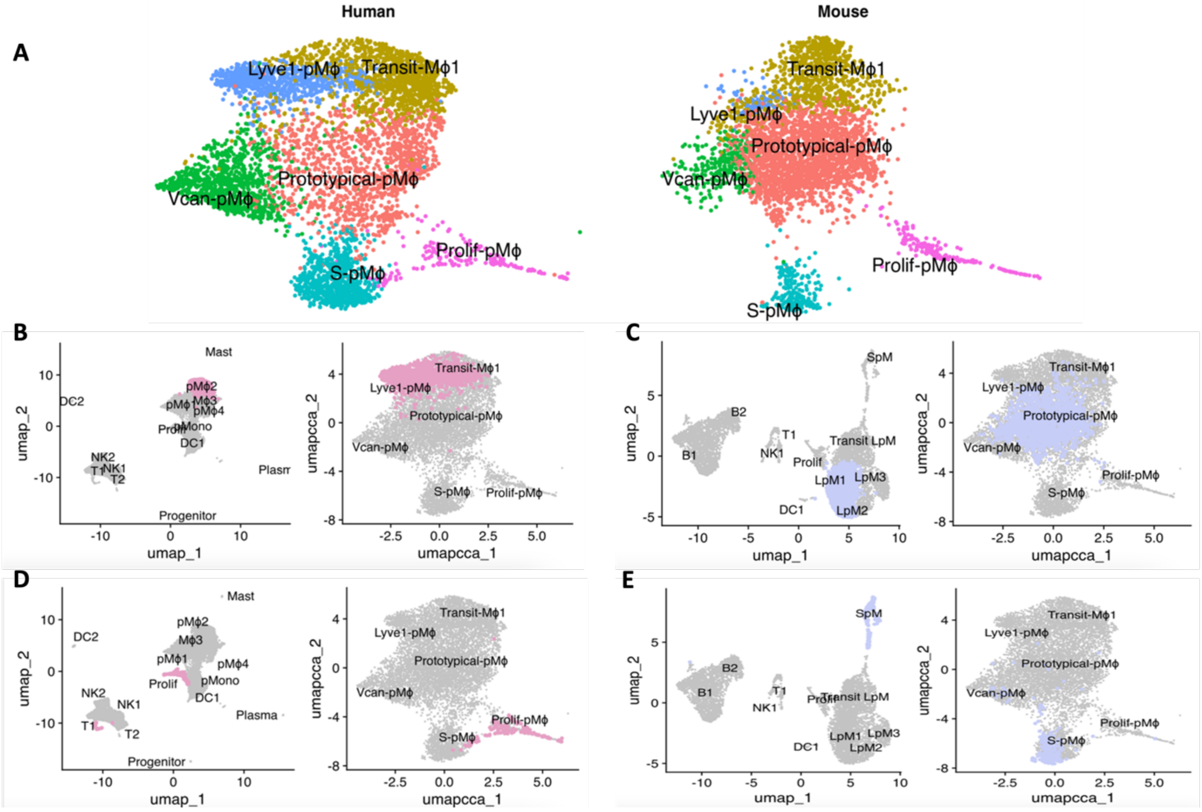


**Figure S6: Validation of integrative clustering.** A) Integrated UMAP separated by species. B) Human pM2 cells mapping to Lyve+ pM and Transit pM clusters in the integrated map. C) Mouse LpM1 (prototypical) cells mapping to the prototypical pM cluster in the integrated map. D) Human proliferating pM cells mapping to the corresponding prolif pM cluster in the integrated UMAP. E) Mouse SpM cells mapping to integrated (S) pM integrated cluster.

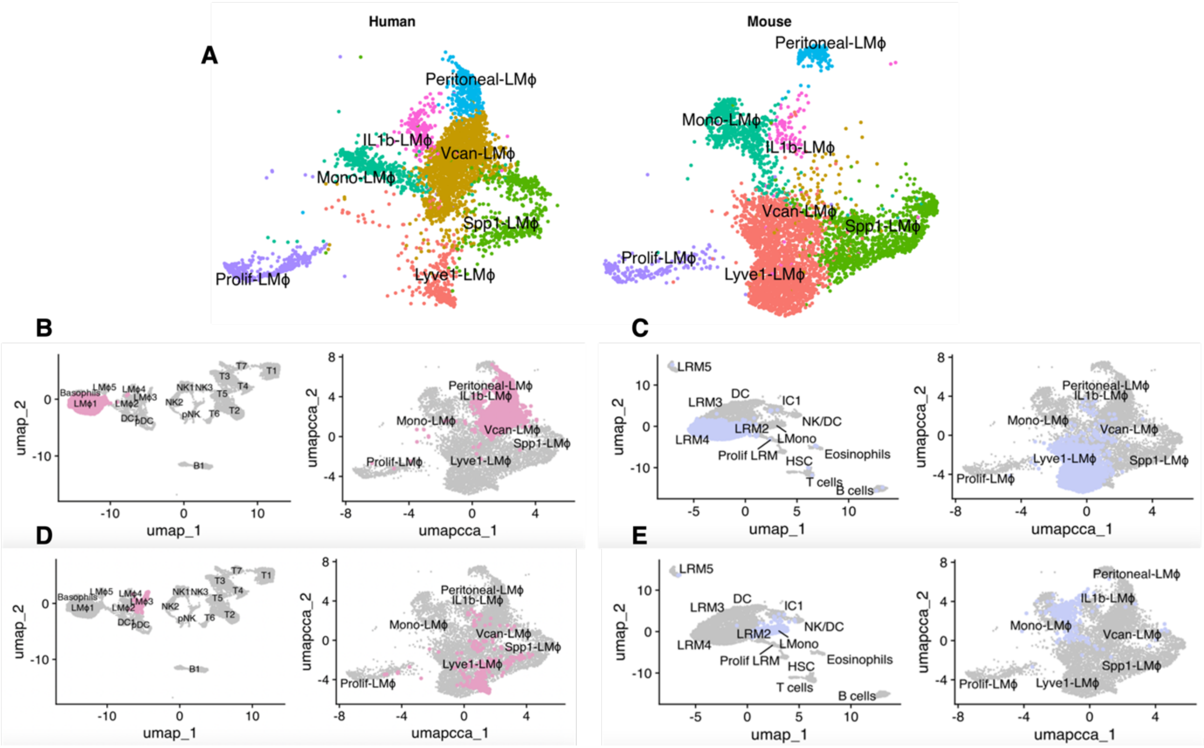


**Figure S7: Validation of integrative clustering.** A) Integrated UMAP separated by species. B) Human LM1 cells mapping to the Peritoneal LM and Vcan+ LM clusters in the integrated map. C) Mouse LRM4 cells mapping to the Lyve1 cluster in the integrated map. D) Human LM3 cells mapping diffusely onto Vcan+ LM, Lyve+ LM and Spp1+ LMin the integrated UMAP. E) Mouse LMono cells mapping to Lmono integrated cluster.
