## Supplementary Table 9 for "Single-cell analysis identifies distinct macrophage phenotypes associated with pro-disease and pro-resolving functions in the endometriotic niche"

| Antibody | **Fluorochrome** | **Dilution** | **Cell Type** | **Supplier + Product Code** | **Clone** | **Application** |
| --- | --- | --- | --- | --- | --- | --- |
| CD3 | APC | 1:300 | T cells | Biolegend 100235 | 17A2 | GFP+/GFP- sorting from MacGreen recipient mice |
| CD19 | APC | 1:300 | B cells | Biolegend 152410 | 1D3/CD19 |  |
| CD335 | APC | 1:300 | NK cells | Biolegend 137607 | 29A1.4 |  |
| Ly6G | APC | 1:300 | Neutrophils | Biolegend 127613 | 1A8 |  |
| Siglec F | APC | 1:300 | Eosinophils | Biolegend 155507 | S17007L |  |
| CD45 | PerCP Cy5.5 | 1:200 | Leukocytes | Biolegend 103132 | 30-F11 |  |
| F4/80 | PE-Cy7 | 1:200 | LpMs | Biolegend 123113 | BM8 |  |
| CD3 | AF700 | 1:300 | T cells | Biolegend 100215 | 17A2 | Analysis of peritoneal fluid macrophages from Apoe mimetic treated mice |
| CD19 | AF700 | 1:300 | B cells | Biolegend 152413 | 1D3/CD19 |  |
| Nk1.1 | AF700 | 1:300 | NK cells | Biolegend 156511 | S17016D |  |
| Ly6G | AF700 | 1:300 | Neutrophils | Biolegend 127621 | 1A8 |  |
| Siglec F | AF700 | 1:300 | Eosinophils | Biolegend 155533 | S17007L |  |
| CD45 | Pacific Blue | 1:100 | Leukocytes | Biolegend 103125 | 30-F11 |  |
| F4/80 | PE-Cy7 | 1:100 | LpMs | Biolegend 123113 | BM8 |  |
| MHC II | PE-Cy5 | 1:100 | SpMs | Biolegend 107611 | M5/114.15.2 |  |
| Timd4 | PE | 1:200 | Tissue resident LpMs | Biolegend 130005 | RMT4-54 |  |
| Ly6C | APC-Cy7 | 1:100 | Monocytes | Biolegend 128025 | HK1.4 |  |
| MHC II | APC-Cy7 | 1:200 | SpMs | Biolegend 107627 | M5/114.15.2 | Sorting of SAM and TAM-like macrophages |
| CD3 | AF700 | 1:300 | T cells | Biolegend 100215 | 17A2 |  |
| CD19 | AF700 | 1:300 | B cells | Biolegend 152413 | 1D3/CD19 |  |
| Nk1.1 | AF700 | 1:300 | NK cells | Biolegend 156511 | S17016D |  |
| Ly6G | AF700 | 1:300 | Neutrophils | Biolegend 127621 | 1A8 |  |
| Siglec F | AF700 | 1:300 | Eosinophils | Thermo Fisher 56-1702-80 | 1RNM44N |  |
| Siglec F | PE/Dazzle | 1:200 | Eosinophils | Biolegend 155529 | S17007L |  |
| FOLR2 | APC | 1:200 | TAMs | Biolegend 153305 | 10/FR2 |  |
| F4/80 | PE-Cy7 | 1:100 | LpMs | Biolegend 123113 | BM8 |  |
| CD206 | PE | 1:100 | TAMs | BD Biosciences 568273 | Y17-505 |  |
| TREM2 | FITC | 1:30 | SAMs | Abcam 119852 | YB2/0 |  |
| CD11b | e-Fluor 506 | 1:200 | Macrophage/Monocyte | eBioscience 69-0112-80 | M1/70 |  |
| CD9 | Vio-blue | 1:50 | SAMs | Miltenyi 130-102-745 | MZ3 |  |
| F4/80 | PE-Cy7 | 1:200 | LpMs | Biolegend 123113 | BM8 | Sorting macrophages for phagocytosis assays |
| MHC II | PE-Cy5 | 1:200 | SpMs | Biolegend 107611 | M5/114.15.2 |  |
| CD45 | PerCP-Cy5.5 | 1:100 | Leukocytes | Biolegend 103132 | 30-F11 |  |
| Timd4 | PE-Cy7 | 1:200 | Tissue resident LpMs | Biolegend 130009 | RMT4-54 |  |
| CD3 | FITC | 1:300 | T cells | Biolegend 100203 | 17A2 |  |
| CD19 | FITC | 1:300 | B cells | Biolegend 152403 | 1D3/CD19 |  |
| CD335 | FITC | 1:300 | NK Cells | Biolegend 137605 | 29A1.4 |  |
| Ly6G | FITC | 1:300 | Neutrophils | Biolegend 127605 | 1A8 |  |
| Siglec F | FITC | 1:300 | Eosinophils | Biolegend 155503 | S17007L |  |
| CD45 | Pacific Blue | 1:100 | Leukocytes | Biolegend 103125 | 30-F11 | Sorting CD45+ cells for single cell sequencing |
| Zombie UV | - | 1:100 | Live/dead marker | Biolegend 423107 | - | Various applications |
| Fixable Viability stain 660 | - | 1:100 | Live/dead marker | BD Biosciences 564405 | - | Various applications |
| Fixable Viability stain 780 | - | 1:100 | Live/dead marker | BD Biosciences  565388 | - | Various applications |
| CD11b | PECF-594 | 1:300 | Macrophage/Monocyte | Biolegend 101256 | M1/70 | Various applications |
